# Genome-wide increased copy number is associated with emergence of super-fit clones of the Irish potato famine pathogen *Phytophthora infestans*

**DOI:** 10.1101/633701

**Authors:** Brian J. Knaus, Javier F. Tabima, Shankar K. Shakya, Howard S. Judelson, Niklaus J. Grünwald

## Abstract

The plant pathogen that caused the Irish potato famine, *Phytophthora infestans*, continues to reemerge globally. These modern epidemics are caused by clonally reproducing lineages. In contrast, a sexual mode of reproduction is observed at its center of origin in Mexico. We conducted a comparative genomic analysis of 47 high coverage genomes to infer changes in genic copy number. We included samples from sexual populations at the center of origin as well as several dominant clonal lineages sampled worldwide. We conclude that sexual populations at the center of origin are diploid as was the lineage that caused the famine, while modern clonal lineages showed increased copy number (3x). Copy number variation (CNV) was found genome-wide and did not to adhere to the two-speed genome hypothesis. Although previously reported, tetraploidy was not found in any of the genomes evaluated. We propose a model of super-fit clone emergence supported by the epidemiological record (e.g., EU_13_A2, US-11, US-23) whereby higher copy number provides fitness leading to replacement of prior clonal lineages.

## Introduction

The Irish famine pathogen, *Phytophthora infestans* (Mont.) de Bary, notorious for destroying the potato crop in Ireland in the 19^th^ century, continues to reemerge globally as one of the world’s costliest plant pathogens (Fry et al. 2015). This pathogen causes late blight on potato worldwide and is considered the most economically important pathogen of this crop. This pathogen is thought to have originated in central Mexico (Goss et al. 2014; Grünwald and Flier 2005) where it is found existing alongside two closely related, endemic sister-taxa defining *Phytophthora* clade 1c, namely *P. mirabilis* and *P. ipomoeae* (Galindo and Hohl 1985; Flier et al. 2002)(Figure 1A). Elsewhere in the world it emerges as clonal lineages (Fry et al. 1993; Hu et al. 2012; Fry et al. 2013; Cooke et al. 2012). These emergent clonal lineages are frequently ephemeral, disappearing after a season or two (Figure S1). However, high fitness (hereafter referred to as ‘super-fit’) clones occasionally emerge and become dominant, replacing the formerly dominant lineages. While this pathogen continues to reemerge globally, we know very little about the mechanisms involved in pathogen emergence and genomic features that are associated with these newly emerging, dominant and super-fit clones.

**Figure 1.**
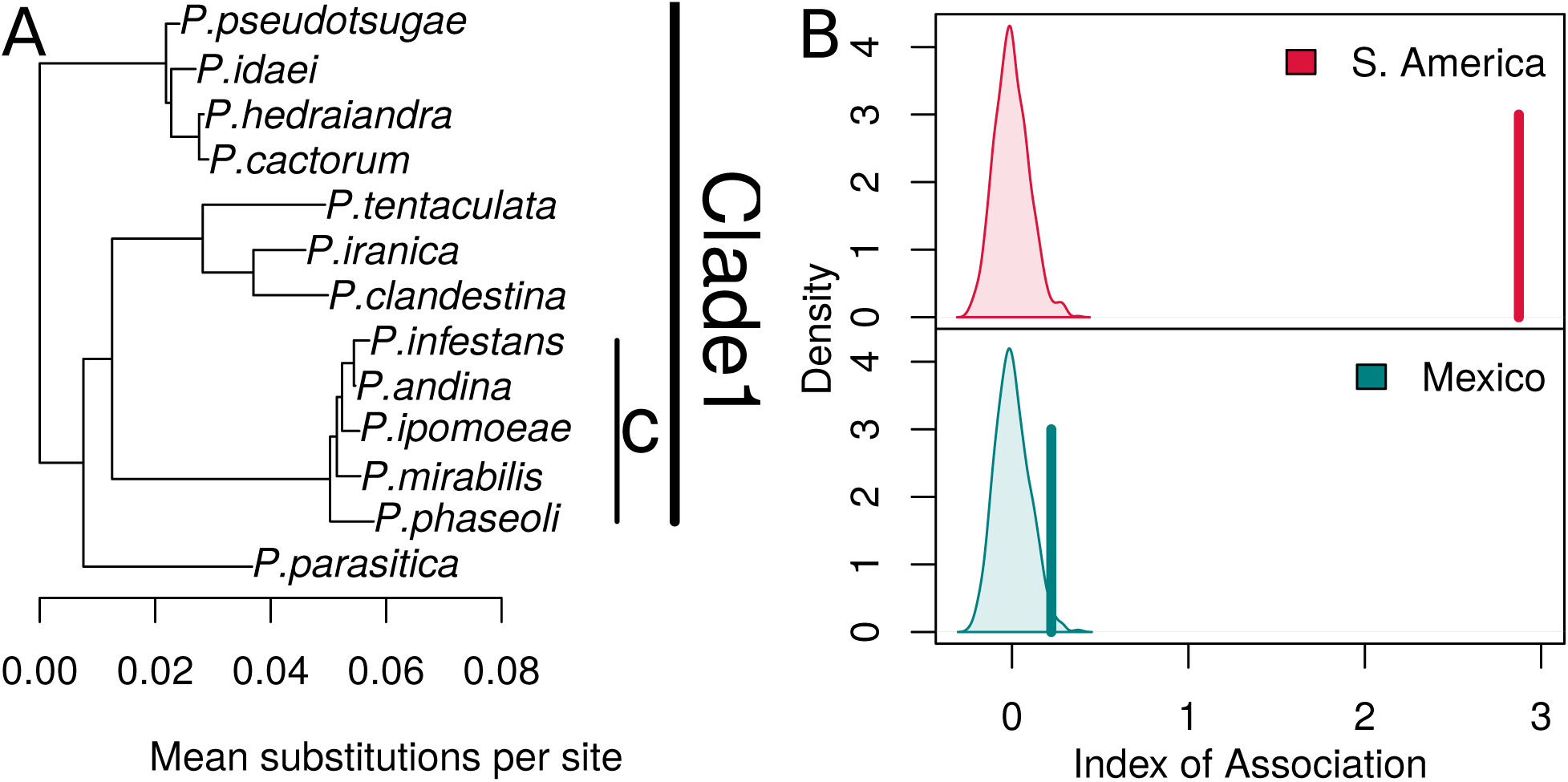
Natural history of the Great Famine pathogen. A. The pathogen *Phytophthora infestans* is considered a member of *Phytophthora* clade 1c (Blair et al. 2008; Kroon et al. 2012; Martin et al. 2014; Flier et al. 2002). The phylogeny is modified from (Martin et al. 2014). B. Populations in Mexico exhibit a sexual mode of reproduction while populations in South America (and elsewhere in the world; data not shown) are generally clonal, or mitotically reproducing (Goss et al. 2014; Flier et al. 2003). Shown are the distribution of simulated expected index of association (*I_A_*) values as well as the observed value (vertical line). In Mexico genetic markers are under linkage equilibrium (P = 0.25); the hypothesis of no linkage among markers is rejected for South American populations (P < 0.001) indicating a clonal mode of reproduction (adapted from Goss et al. 2014).

*P. infestans* exhibits two distinct lifestyles worldwide. At the center of origin in central Mexico, the pathogen exists as a sexual, randomly mating population (Goss et al. 2014; Grünwald and Flier 2005) (Figure 1B). Throughout much of the remainder of the world, *P. infestans* is distributed as distinct clonal lineages that reproduce mitotically. Until the early 1990s a single lineage, US-1, dominated the global populations (Goodwin et al. 1994). US-1 was thought to be the lineage that caused the Great Famine. However, more recent work identified HERB-1 as the famine causing lineage (Yoshida et al. 2013), a lineage that differs from US-1, but might be ancestral to US-1 (Martin et al. 2013, 2016). During the mid-1990s, late blight re-emerged in the US as novel clonal genotypes that were not previously observed (Fry and Goodwin 1997a, 1997b). The epidemiologically most notable genotypes included US-8 and US-11 that were characterized as having resistance to the fungicide metalaxyl. During the late 2000s novel lineages emerged in the US including US-22, US-23, and US-24 (Hu et al. 2012; Fry et al. 2013). Similar observations were made in Europe where the 13_A2 clonal lineage became dominant in the late 2000s where it displaced 6_A1 in the UK and other previously existing clonal lineages (Cooke et al. 2012). The global population structure of *P. infestans* is therefore characterized as having a sexually reproducing population at its center of origin as well as re-emerging clonal epidemics in the US, and most of the rest of the world, consisting of distinct clonal lineages that displace older clonal lineages.

The *P. infestans* genome has been characterized as having a two-speed genome. These two speeds refer to two compartments: gene-dense regions, containing predominantly housekeeping genes, and gene-sparse regions enriched for effectors (proteins that are secreted from the pathogen and associated with infection) including RxLR genes (Haas et al. 2009; Raffaele et al. 2010). It is thought that dramatic changes to the gene-sparse, transposon and effector rich portion of the genome are responsible for most of the adaptation in clonal lineages. For example, Cooke et al. (2012) studied the recent emergence of the 13_A2 clonal lineage in the UK that largely displaced clonal lineages existing in the UK by about 2008. This study documented that this lineage was more aggressive, thus out-competing and displacing older lineages. They also reported large changes in copy number variation (CNV), gene loss, mutations, and gene expression patterns that distinguished 13_A2 from previous lineages. These genomic changes are thought to underlie its emergence.

In addition to the two-speed genome model, several studies have documented variation in ploidy. *Phytophthora* species are considered to be diploid (Brasier 1992). Extensive cytological work documented that *P. infestans* was primarily diploid, yet indicated that some isolates might be of higher ploidy (Sansome and Brasier 1973; Sansome 1977). Several cytological studies indicated that individuals from sexual populations in Mexico were diploid, whereas individuals from clonal populations elsewhere frequently exhibited higher levels of ploidy (Tooley and Therrien 1987; Catal et al. 2010). More recently, Yoshida et al. (2013) analyzed whole genome sequences to show that the allele balance (e.g., the frequency of each allele sequenced at heterozygous positions) for some individuals was triploid or tetraploid. This observation of higher ploidy was further supported by work combining high throughput sequencing and flow cytometry (Li et al. 2017). This body of prior cytological and genomic work provides support for a model that clonal populations are often triploid or tetraploid while some populations/strains might be diploid. However, these observations are based on individual samples not allowing broader inferences about populations at large and have not included a representative sample from sexual populations.

We re-sequenced genomes of *P. infestans* to explore variation in gene copy number and in a representative global sample that included a sexual population and select members of clonal lineages. We combined our genome data with recently published whole genome data to obtain a population of 47 high coverage samples (Figure 2) that provide power for testing the hypotheses of finding difference in ploidy, CNV and genic content in *P. infestans.* For this study, we defined ploidy as a genome wide change in copy number (i.e., whole genome duplication), whereas copy number refers to a change observed at the sub-chromosomal level. We tested the hypotheses that sexual populations were diploid with little CNV while clonal populations were predominantly triploid or tetraploid with high CNV. We also tested the hypothesis that CNV and presence absence polymorphism are enriched in gene-sparse, effector rich portions of the genome (as expected by the two-speed genome hypothesis). We also expected to find CNV and presence/absence polymorphisms differed in clonal vs. sexual populations. Finally, we tested the hypothesis that similar changes in CNV might be observed in other heterothallic *Phytophthora* species for which genomic data for populations was available, such as *P. parasitica* and *P. capsici*. Our findings provide a new perspective on how plasticity in ploidy, copy number and presence/absence polymorphisms contribute to the emergence of the Irish potato famine pathogen and other *Phytophthora* pathogens.

**Figure 2.**
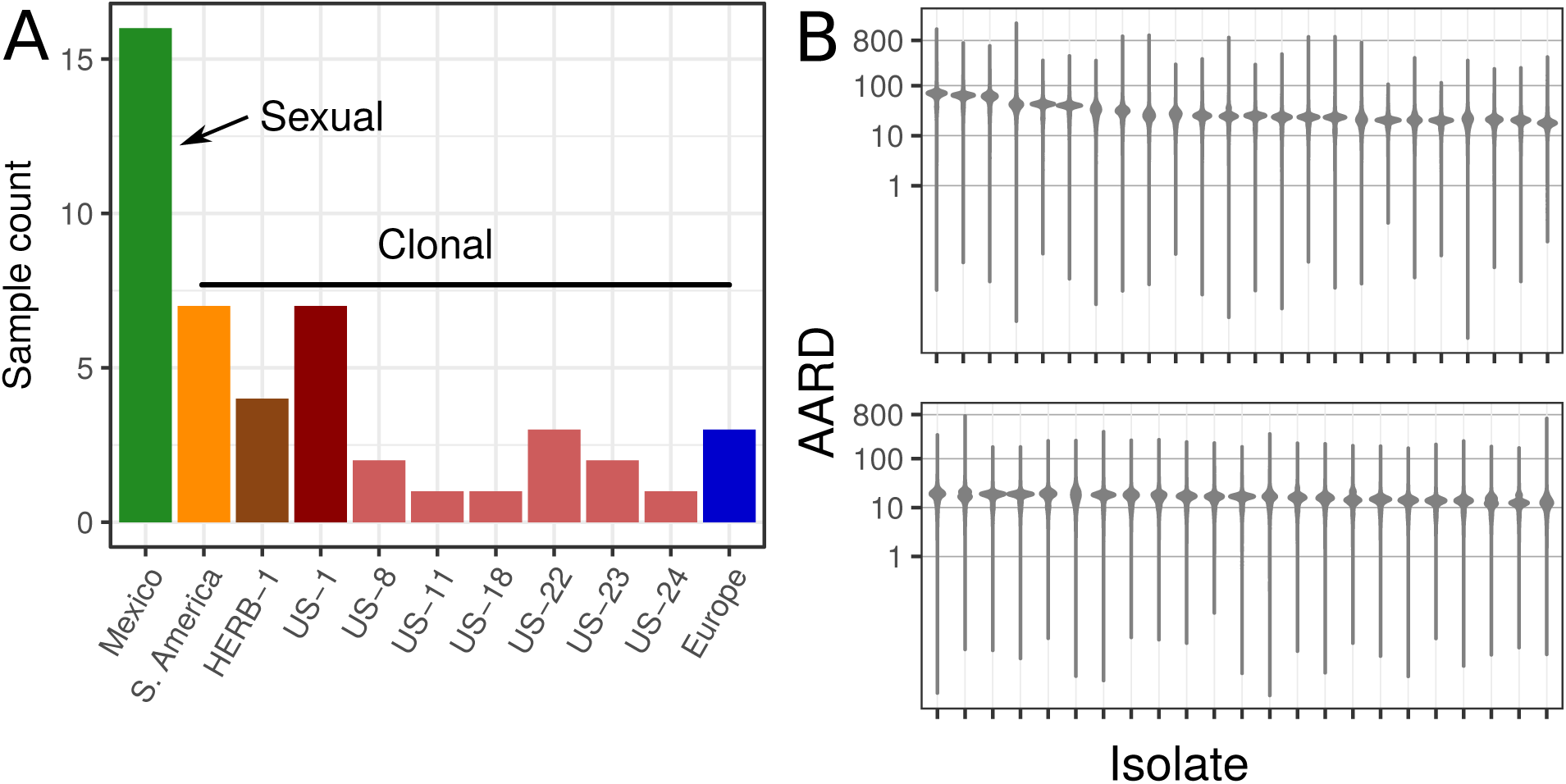
Samples included sexual and clonal populations selected for high sequence coverage. A. The sample was collected from throughout the world and included samples from Mexico that are sexually reproducing (green bar) as well as samples from the rest of the world that are clonally reproducing (all other bars). B. In order to attain high quality samples from the literature and our own resequencing for the inference of copy number variation, only samples with at least 12X adjusted average read depth (AARD) were included.

## Results

### Resequencing populations of *P. infestans*

To understand variation in CNV and gene content, we resequenced and used previously published, populations of the potato late blight pathogen, *P. infestans*, from the center of origin in Mexico (n = 16), and dominant clonal lineages in the US, Europe and South America (Figure 2A). To allow for robust inference of gene copy number we used only genomes with a genic average adjusted read depth (AARD) of 12x or greater (Figure 2A; Figures supplement 2A, B, C). This resulted in a total of 47 high quality *P. infestans* genomes (Supplementary table 1; Figure 2B).

### Genic copy number varies continuously in *P. infestans*

We observed genic CNV among populations (Figure 3A) and a gradient of genic copy number ranging from predominantly 2x to predominantly 3x (Figure 3B). We did not observe classes of individuals that would represent tetraploid individuals. Isolates from the United States belonging to clonal lineages have a gradient of gene copy number (Figure 3C). Diploid strains in US lineages were mostly found in the well-represented lineage US-22 (n = 3) and US-23 (n = 1). Similarly, in Europe isolates that were both predominantly diploid and triploid were observed. The exception to this balance of ploidy appeared to be in South America where almost the entire sample appeared to be predominantly triploid (Figures 3C). Gene copy number for samples from Mexico demonstrated a greater number of strains that had genes with predominantly two copies. While previous studies have focused on variation in ploidy (Sansome and Brasier 1973; Sansome 1977; Tooley and Therrien 1987; Catal et al. 2010; Yoshida et al. 2013; Li et al. 2017) our work supports variation in genome size in *P. infestans* occurring largely at a sub-genomic level: Mexican, Herb-1 and US-22 samples were predominantly 2x with narrow variation that can be interpreted as diploidy, whereas samples from South America, US-1, other US lineages and Europe showed large variation (Figure 3A).

**Figure 3.**
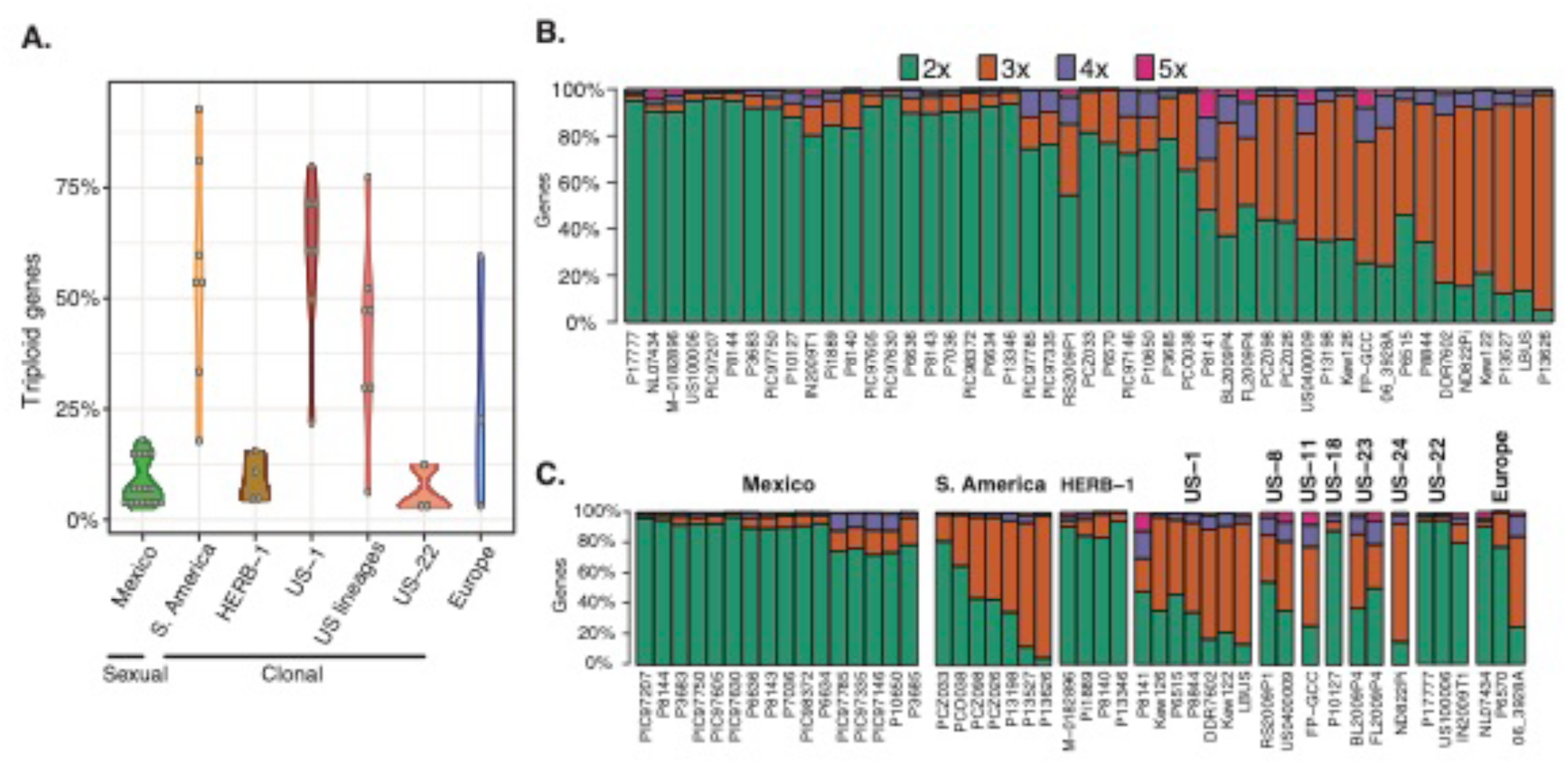
Strains of *P. infestans* show variation in gene copy number. A. Populations from Mexico, HERB-1 and US-22 are predominantly diploid while populations from elsewhere in the globe are predominantly triploid. **B.** Barplots of the proportion of inferred gene copy number based on allele balance over all genes for each individual. The top row shows the entire population showing a gradient in the proportion of gene copy number across all samples. **C.** The second bar chart organizes the first bar chart by origin starting on the left from Mexico, South America, HERB-1 (the Irish Famine lineage (Yoshida et al. 2013)), US-1, US-8, US-11, US-18, US-22, US-23 (currently most abundant in US; see Figure supplement 1), US-24 and finally, Europe.

The degree of CNV was also explored for samples where tissue was extracted from historical herbarium samples (HERB-1: M-0182896, Pi1889; US-1: Kew122, Kew126) (Figure 3C). These samples were not cultured on media and were not exposed to the modern fungicide metalaxyl. These samples demonstrated variability in gene copy number as well (Figure 3A, C) suggesting that CNV may have been a natural condition in clonal lineages of *P. infestans*. Note that two of the four samples that we determined to be of sufficient sequence depth to call copy number were from the 20^th^ century (Kew122 and Kew126 both collected in 1955) and clustered with US-1 (Martin et al. 2016) while the other two are from the 19^th^ century and clustered with Herb-1 (M-0182896 collected in 1877 and Pi1889 collected in 1889). This indicates that CNV was observed throughout the time series of the data and was not restricted to modern samples that were cultured on media.

### Gene loss occurs in both clonal or sexual populations

We explored the hypothesis that gene loss (relative to the reference genome T30-4) had occurred collectively within a lineage or independently. The breadth of coverage (BOC) for a gene is the proportion of positions that were sequenced at least once in the reference genome (Raffaele et al. 2010). For example, a BOC of 0.75 would indicate that 75% of the positions in a gene were sequenced at least once. We used a BOC of zero to define a gene loss event and presented samples for populations that included at least six individuals (groupings with more were randomly subset to a sample size of six)(Figure 4; Figure supplement 5). Gene loss was most pronounced in RxLR and CRN effectors but was found in all gene classes (Figure supplement 5). Gene loss among the isolates from Mexico ranged from 38 to 112 gene deletions. However, we found only one shared deletion among all samples within the clonal lineage (Figure 4, bottom panel). Clonally reproducing isolates from South America demonstrated a loss of 39 to 63 genes with only 9 gene losses shared in common among these isolates. Among 6 individuals belonging to lineage US-1 we observed a range of loss from 21 to 68 genes, but only 5 gene losses common among all of the sampled lineages (Figure 4). Gene loss is a common feature in the gene sparse regions of the genome. However, the specific gene lost within any particular sample is unique and random and apparently affects clonal and sexual populations equally.

**Figure 4.**
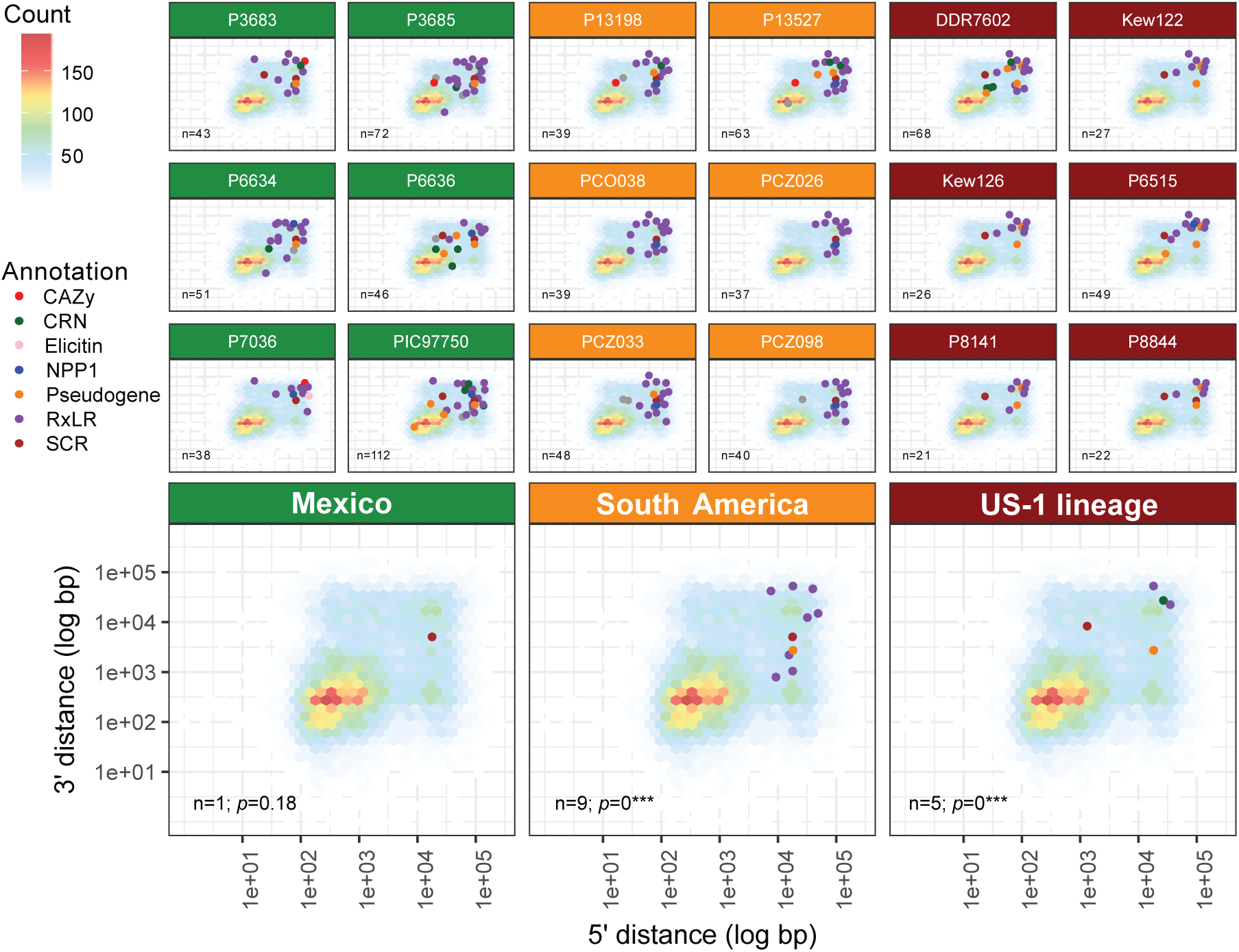
Gene loss in pathogenicity factors and pseudogenes occurs in individuals within populations regardless of clonal or sexual mode. The top three rows show results for individual samples and the bottom row shows the summary for the populations sampled from Mexico (green), South America (orange), and the US-1 clonal lineage (red). Gene loss is common in individuals where any individual may lose between 21 and 112 pathogenicity genes or pseudogenes. However, when comparing multiple samples within a population few losses are shared. Gene loss is happening on an individual basis and not dependent on ancestry. This pattern is observed in sexually reproducing populations (Mexico) as well as in clonally reproducing populations (South America and US-1). Gene loss is defined here as a gene from a sample with an adjusted average read depth of at least 12 where zero positions (base pairs) in the gene were sequenced (BOC = 0). The background for each panel is a heatmap indicating gene abundance in relation to their 5’ and 3’ intergenic spacing. Points indicate gene deletions and are colored by their functional annotation as defined in Haas et al. (2009).

### Genic copy number variation was not associated with specific classes of genes

If genic copy number was associated with the phenotype we would expect CNV to be associated with a particular type of gene class (e.g., pathogenicity factors). We found that in the sexually reproducing population from Mexico that was predominantly diploid all gene categories had more 2X genes than 3X genes (Figure 5; green). Similarly, for the populations from S. America (orange) and US-1 (red) that were clonally reproducing we found that all gene classes had more 3X genes than 2X genes regardless of gene family. CNV occurs throughout gene space without a preference for functional annotation (Figure 5).

**Figure 5.**
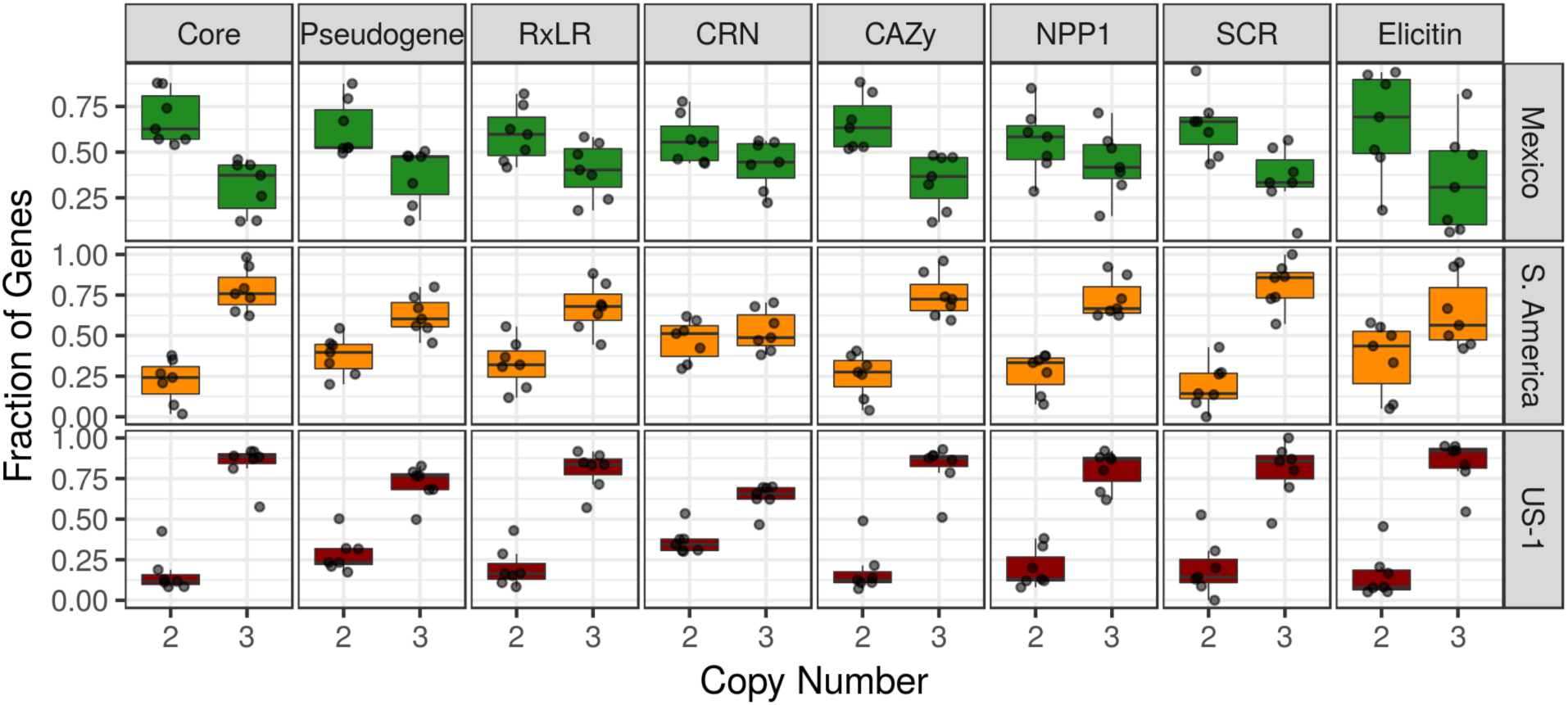
Gene copy number variation is not restricted to a particular class of gene. Columns show gene families and rows show results by population. Isolates from Mexico (green), where *P. infestans* is sexually reproducing, had a gene copy number predominantly of two for all classes of genes. Isolates from South America and US-1, both considered clonally reproducing, had a gene copy number predominantly of three for all gene classes. Gene copy number varies throughout gene space and is not associated with function. Box and whisker plots summarize points that represent samples (n=6) and the proportion of genes that were either 2x or 3x (based on the total number of 2x and 3x genes). A sample size of six was used (as in figure 4) to have equal samples sizes. Core = core orthologous genes; other gene families are defined as in Haas et al. (2009).

### Gene copy number variation occurred in core orthologous genes

Core orthologous *Phytophthora* genes are genes that were reported to occur once and only once in *P. infestans*, *P. ramorum*, and *P. sojae* (Haas et al. 2009) and are thought to be highly conserved. Therefore, we would expect only low levels of CNV in these genes. To test this hypothesis, we plotted all core orthologous genes present at 3x by their 5’ and 3’ intergenic distances (Figure 6). We observed substantial numbers of genes inferred to have three copies (3x) among core orthologous genes in the gene dense portion of each genome (Figure 6). This indicates that this portion of the genome may be more dynamic than previously thought.

**Figure 6.**
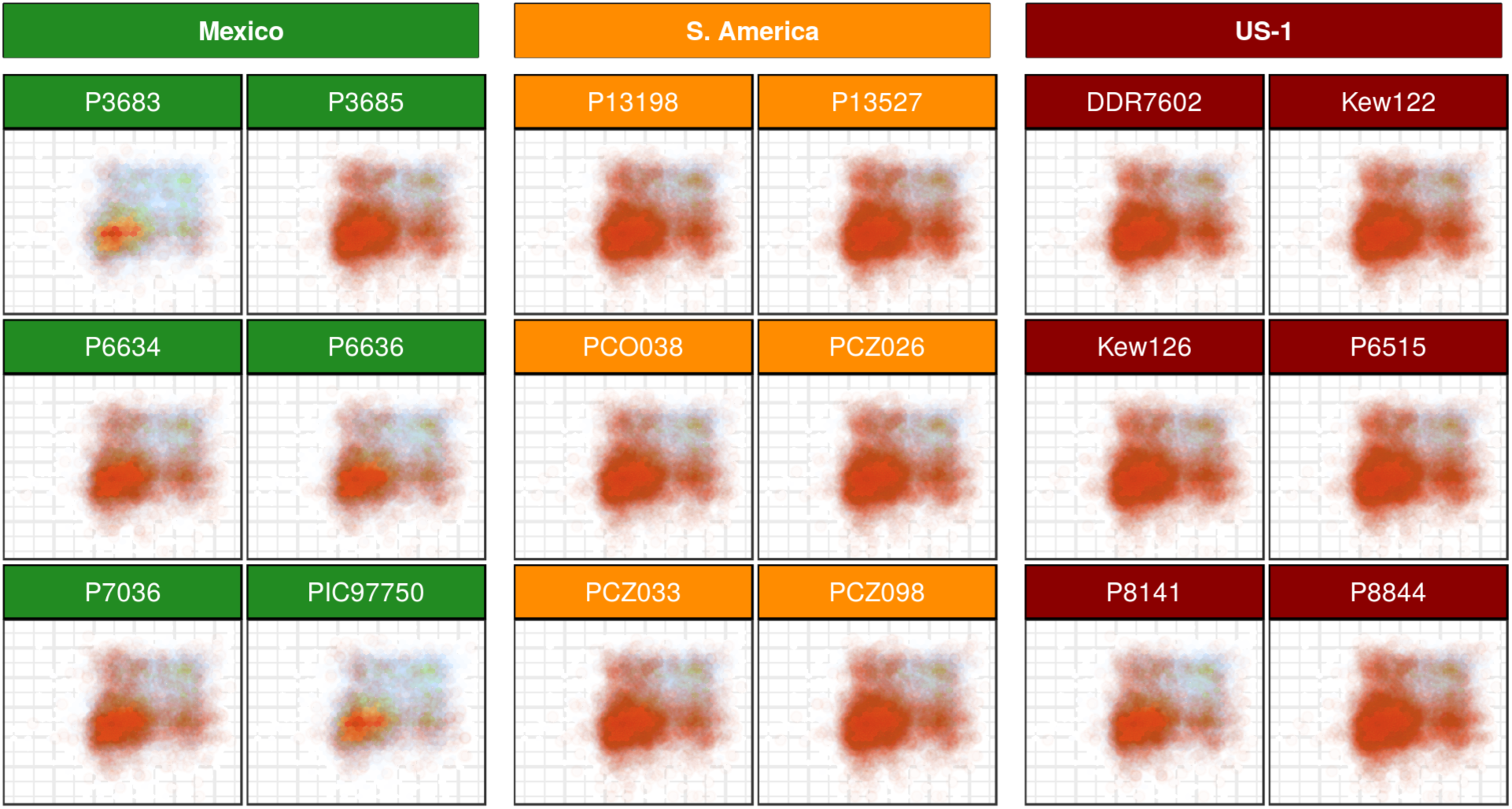
Gene copy number does not follow the two-speed genome hypothesis. Core orthologous genes with a copy number of three are enriched in the gene-dense (rather than gene-sparse) regions of the genome. The background for each panel is a heatmap indicating gene abundance as in Figure 4. Orange points (with transparency) are plotted over this background where each point is a core orthologous gene that was determined to have three copies and was positioned based on their 3’ (y-axis) and 5’ (x-axis) intergenic distance as in Figure 4.

### The phenomenon of genic CNV is shared with other members of the *Phytophthora* genus

We explored if variation in ploidy apparent in *P. infestans* is observed in other heterothallic *Phytophthora* taxa. We looked at species for which population level genome data was available including *P. andina* (clade 1c), *P. parasitica* (clade 1), and *P. capsici* (clade 2)(clades as assigned by Blair et al. 2008). The taxon *P. andina* appears to be diploid in our limited sample (Figure 6). However, we observed more heterozygous positions than in the other taxa (Figure 6). This is consistent with the interpretation that *P. andina* is a homoploid hybrid that arose from a cross between *P. infestans* and another undescribed *Phytophthora* species (Goss et al. 2011). The more distantly related *P. parasitica* appeared diploid as well. However, its relatively high sequence depth allowed resolution of minor peaks indicating that a fraction of genes occur at three copies (particularly in the sample P1569). The taxon most distantly related to *P. infestans* included in our analysis was *P. capsici*. Three of the *P. capsici* samples appeared to be diploid while one sample (Pc389) appeared to be triploid. These results suggest that our findings of variation in ploidy and CNV within *P. infestans* are also shared among other species of *Phytophthora*.

**Figure 7.**
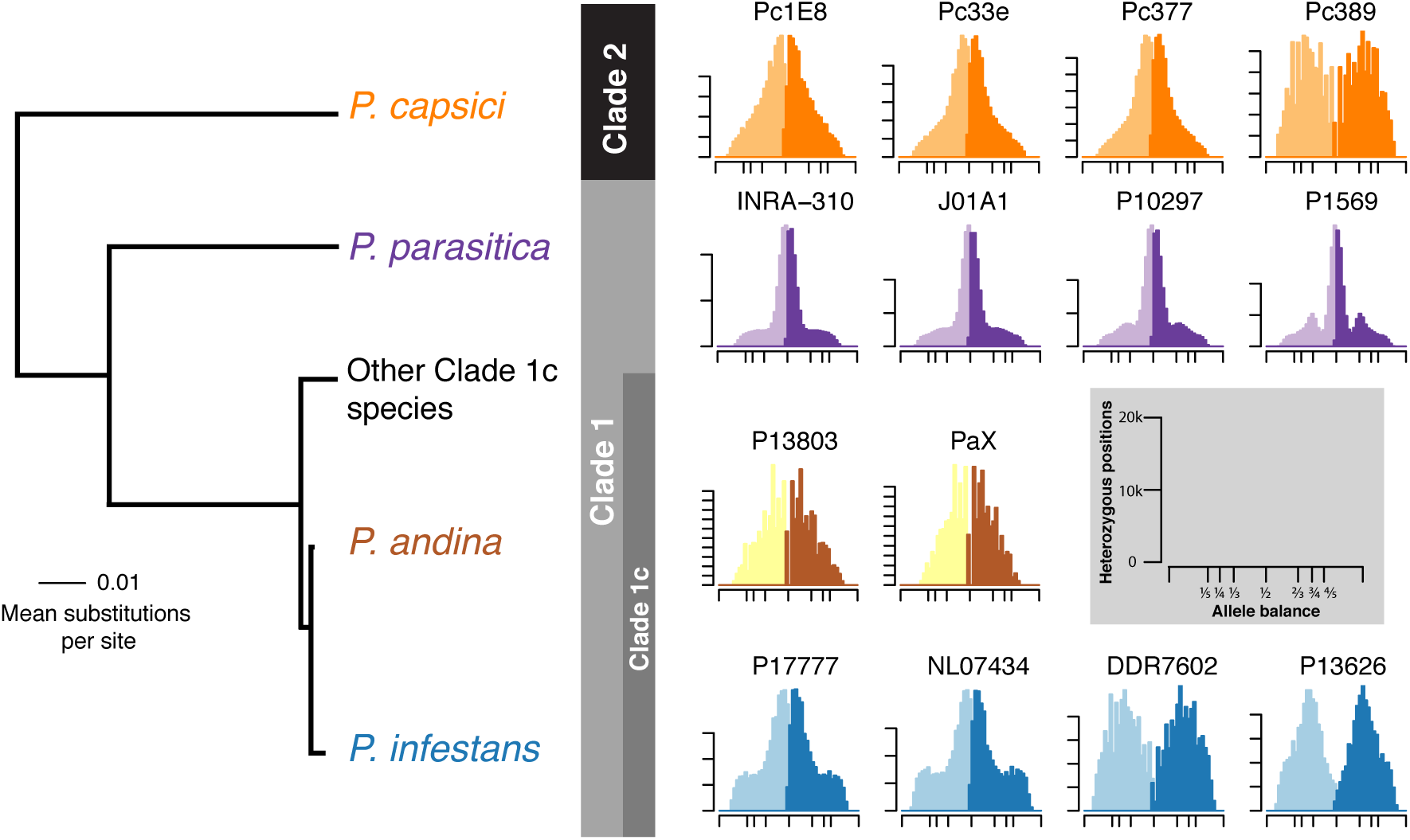
Ploidy variation is observed in other heterothallic *Phytophthora* species. Histograms showing the major and minor allele frequency observed for individuals. Samples of *P. infestans* are diploid or triploid. The hybrid taxon *P. andina* appeared predominantly diploid. Note that the y-axis indicates many more heterozygous positions in these samples relative to the other species. The clade 1 species *P. parasitica* is diploid. However, note the ‘shoulder’ peaks at the 3X expectation for samples P10297 and P1569. The clade 2 species *P. capsici* demonstrated a predominance of diploidy with a single sample being triploid. The x-axis ticks mark 0, 1/5, 1/4, 1/3, 1/2, 2/3, 3/4, 4/5 and 1 marking our expectations for pentaploid, tetraploid, triploid and diploid. The y-axis ticks in the histograms are in units of 10,000 heterozygous sites. Phylogeny adapted from (Martin et al. 2014). Larger histograms are presented in Figure supplements 3-4.

## Discussion

To characterize the emergence of new clonal lineages of the Irish famine pathogen *Phytophthora infestans* we resequenced whole genomes of select populations from several dominant clonal lineages as well as the first sexual populations from the center of origin in Mexico. Prior work focused primarily on select individuals rather than populations and did not include sexual populations. The genomes were compared with previously sequenced, high quality genomes to determine ploidy, CNV and gene content. Recent epidemiological records indicated that new clonal lineages have emerged repeatedly in the US and Europe (Figure supplement 1). For example, the lineage US-1 was the first to establish in the US, but was eventually displaced by US-8, US-11 and more recently by US-23 (Fry et al. 2015)(Figure supplement 1). Similarly, populations in the UK were displaced by 13_A2 in the last decade and more recently 6_A1 (Cooke et al. 2012). While variation in ploidy has been described in select individuals from clonal lineages of *P. infestans* our work provides several new key insights based on population-level patterns expanding on prior work focusing on single clonal strains.

### Clonal lineages show higher copy number than sexual populations at the center of origin

Populations studied show a gradient of CNV from 2x to 3x (Figure 3B). Populations of *P. infestans* that are sexually reproducing at the species’ center of diversity in Mexico are diploid (Figure 3C). This contrasted with dominant clonal populations from the rest of the world that are predominantly triploid. This suggests that there may be a connection between ploidy, epidemic fitness, and mode of reproduction.

### Isolates were predominantly diploid or triploid, but not tetraploid

We observed no tetraploid individuals as reported previously (Yoshida et al. 2013). In fact, we reanalyzed some of the same samples and data including the European lineage 13_A2 previously characterized as being tetraploid. In our analysis 13_A2 had mostly three gene copies (Figure 3), which is in agreement with a more recent report (Li et al. 2017). Part of this discrepancy is due to changes in technology. Plotting histograms of allele balance has typically included all variants, including homozygous genotypes. Because homozygous sites are much more abundant than heterozygous sites this tends to drive the scaling of the plot. To avoid this, previous work limited plots to a frequency range of 0.2 to 0.8. We subset our data to only the heterozygous genotypes, resulting in a plot from 0 to 1, and subset the data by omitting variants with unusually high or low sequence depth. This is a significant improvement in methodology for inferring ploidy or CNV based on allele balance (Knaus and Grünwald 2018).

### Gene loss occurred within individuals in both sexual populations and clonal lineages

We tested the hypothesis that gene loss was shared by ancestry. This would provide the expectation that members of a clonal lineage show fixed polymorphisms within members of the clonal lineage. We used breadth of coverage to identify presence/absence of genes relative to the reference genome. Instead we found that individuals within a clonal lineage (e.g. from South America or US-1; Figure 4) showed gene loss within individuals at a similar rate to the sexual population. Furthermore, gene loss affected many gene families including effectors and was located throughout the genome. This is in line with the hypothesis where pathogenicity factors are thought to be enriched in the gene sparse portion of the genome (Haas et al. 2009; Raffaele and Kamoun 2012; Dong et al. 2015).

### CNV is found throughout the genome and affects all gene families including core genes and effectors equally

Our expectation following the proposed two-speed genome hypothesis was to find CNV enriched in the gene-sparse, transposon and effector rich portion of the genome where CNV could provide a means of creating novel paralogs. To our surprise CNV affects housekeeping genes and effectors equally and is randomly dispersed throughout the whole genome (Figure 5). In the diploid genomes from Mexico we found that core orthologous genes, pseudogenes, and several pathogenicity factors were all predominantly 2x (Figure 5). Genomes of clonally reproducing strains from South America and the lineage US-1 were found to have core orthologous genes, pseudogenes, and pathogenicity factors that were predominantly 3x. We also expected CNV to be higher in pathogenicity factors than in core orthologs, yet levels of CNV were not different regardless of gene class (Figure 5).

### Variation in ploidy can be found in other but not all *Phytophthora species*

We also evaluated if changes in ploidy could be observed in other heterothallic *Phytophthora* species. We used genomes for moderate population sizes from *P. andina*, *P. parasitica* (= *P. nicotianae*), and *P. capsici* available at the sequence read archive to address this question (Figure 6). Within *Phytophthora* clade 1c, *P. andina* appeared predominantly diploid. *P. andina* has been recognized as a hybrid with two parental species, one of which is *P. infestans* while the other hybrid parent is unknown (Goss et al. 2011; Oliva et al. 2010). The genomes of *P. andina* had one haplotype from each parental species as expected and were predominantly 2x copy number. *P. parasitica*, a distant relative of *P. infestans* basal to clade 1, was diploid. However, two strains (P10297 and P1569) had minor peaks at our expectation for three copies indicating that fractions of these genomes may vary in copy number at 3x. Our ability to resolve these peaks was likely due to the high sequence depth of these samples relative to the other available taxa. In clade 2, the more distant *P. capsici* appeared predominantly diploid for 3 strains; however, one strain (Pc389) was triploid. These results suggest that variation in ploidy and/or copy number may be a common feature throughout the *Phytophthora* genus.

### We propose a model of emergence where super-fit triploid clones emerge and eventually displace prior clonal lineages

Our work provides striking support for a model of predominantly diploid populations at the center of origin reinforced by sexuality and predominantly triploid clonal lineages elsewhere in the world (Figure 8). In this model, novel clonal lineages emerging globally are predominantly triploid. These triploid lineages are super-fit and displace other extant lineages. A new lineage emerging from a sexual cross in Mexico is expected to be initially diploid and will gradually show 3x CNV. Older previously dominant lineages might thus be more triploid (e.g., US-1) than dominant younger lineages (e.g., US-23). Some lineages are ephemeral (e.g., US-18, US-22). The recently emerged diploid lineage US-22 was only observed between 2009-11 and might be less fit (Danies et al. 2014; Fry et al. 2015) and curiously shows predominantly 2x CNV. To the best of our knowledge all lineages that became dominant in space and time are or were triploid with the exception of HERB-1. It remains to be established if higher genic copy number confers higher epidemic fitness to a clonal lineage. Experimentally addressing this question might prove challenging given the fact that CNV is a whole-genome phenomenon.

**Figure 8.**
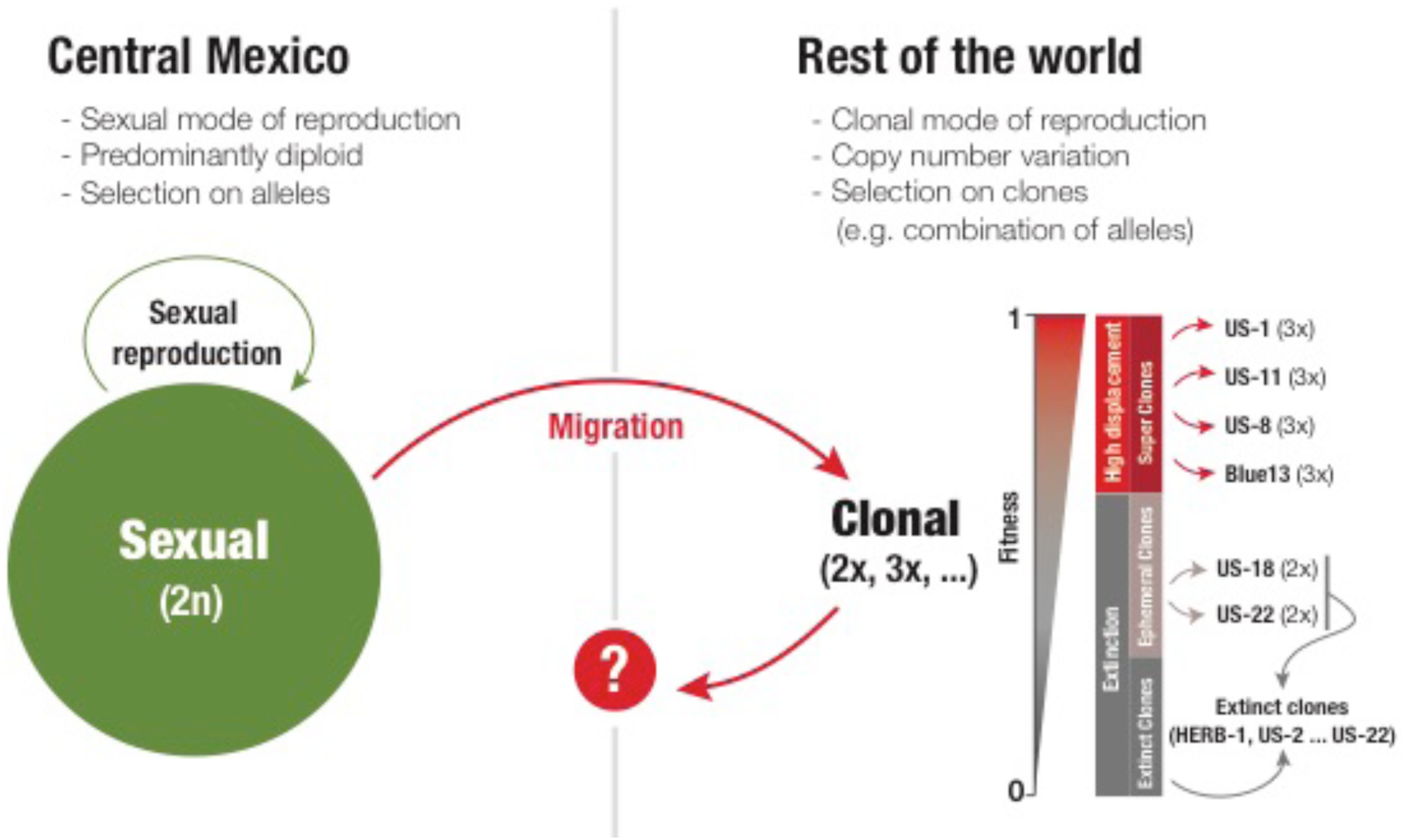
Model of the genomic processes observed to date thought to underlie the patterns of emergence observed for clonal lineages of the Irish famine pathogen. This model proposes that sexual populations are diploid, while clonal populations are predominantly triploid. Some clones, likely rare and by chance, are super-fit clones that become dominant, displacing prior clonal lineages, as has been observed in the US, repeatedly. These super-fit clones are predominantly triploid including US-8, US-11, US-23, and Blue13_A2 while other lineages that are diploid such as US-18 and US-22 are ephemeral or cryptic (Figure supplement 1). Sexual reproduction reinforces diploidy while a triploid status might interfere with sexual reproduction but might confer fitness.

## Conclusions

The late blight pathogen *P. infestans* continues to re-emerge causing financial loss for farmers and threatening food security, particularly in developing countries (Fry et al. 2015). We report the observation that *P. infestans* isolates are diploid in central Mexico where they reproduce sexually and dominant clonal lineages are predominantly triploid. These findings provide novel support for the hypotheses that a change in copy number might drive emergence of clonal lineages of the Irish Famine pathogen.

## Materials and methods

### Sequence alignment and variant calling

The sample came from previously published sources (Raffaele et al. 2010; Haas et al. 2009; Cooke et al. 2012; Yoshida et al. 2013; Martin et al. 2013, 2016) as well as 11 new *Phytophthora infestans* genomes we sequenced (Table S1). Isolates US040009, FP-GCC, US100006, FL2009P4 and ND822Pi were sequenced at the UC Davis Genome Center. Isolates PIC97136, PIC97146, PIC97335, PIC97442 and PIC97750, and PIC97785 were sequenced at Oregon State University’s Center for Genome Research and Biocomputing on an Illumina HiSeq2000. Additionally, five samples each of *P. mirabilis* and *P. ipomoeae* (Table S2) were also sequenced at Oregon State University’s Genome Research and Biocomputing on an Illumina HiSeq2000. All other samples were obtained from publicly available repositories (Supplementary tables 1 and 2). Newly sequenced genomes are publicly available at the Sequence Read Archive (BioProject ID: PRJNA542680; Supplementary table 1).

The FASTQ format files were aligned to the *P. infestans* T30-4 reference (Haas et al. 2009) using bwa mem 0.7.10 (Li and Durbin 2009; Li et al. 2009). The resulting SAM format file was converted to BAM format, had mate information fixed and had the MD and NM tags added using SAMTools (Li et al. 2009). PCR and optical duplicates were marked using Picard’s MarkDuplicates (Broad Institute n.d.). Per gene sequence depth and coverage over all T30-4 genes was calculated using SAMtools mpileup (Li et al. 2009). From the mpileup data, the number of positions that were sequenced at least once and a median of coverage were calculated. In order to correct our measure of coverage for GC bias we calculated an adjusted average read depth (AARD) (Raffaele et al. 2010). A median was chosen as a robust alternative to an average, however we refer to our measure here as AARD to be consistent with the existing literature. The genes were sorted into bins based on percentiles of GC content. The adjusted median read depth was then taken by multiplying the median read depth for each gene by the ratio of the median read depth of all genes divided by the median average read depth for all genes in the GC bin of the gene. The AARD for each genome was summarized using violin plots (Wickham 2009) and a threshold of mean AARD of at least 12 was used as a threshold for inclusion of a genome for further analysis.

Variants were called from the BAM files for diploid genotypes to create genomic variant call format (gVCF) files using the Genome Analysis Toolkit (GATK) HaplotypeCaller (Auwera et al. 2013; DePristo et al. 2011). Diploid genotypes were called using the GATK’s GenotypeGVCFs. The samples P10127, P10650, P11633, P12204, P1362, P6096 and P7722 were flagged by the GATK’s HaplotyeCaller as having legacy quality encoding. These samples were run with the option fix_misencoded_quality_scores to accommodate this.

### Gene copy number inference based on allele balance

Inference of gene copy number was made based on the ratio of alleles observed at heterozygous positions (Yoshida et al. 2013; Li et al. 2017). The VCF specification (Danecek et al. 2011) provides the option for variant callers to report the number of times each allele was sequenced at a variable position. In a diploid heterozygote the expectation is that each allele will be observed at an equal frequency, or a ratio of one half. A triploid heterozygote will be expected to have alleles observed at a ratio of one third. A tetraploid heterozygote will be expected to have alleles observed at a ratio of one quarter. Note that some combinations are indistinguishable and therefore uninformative. For example, a tetraploid heterozygote with only two alleles (e.g., A/A/C/C) will have each allele observed at a ratio of one half. This will be indistinguishable from our expectation from a diploid heterozygote. The ratio of alleles observed at each variable position has been used by other authors to make inferences about ploidy (Yoshida et al. 2013; Li et al. 2017). Shortcomings of the present use of the ratio of alleles are that it has been presented graphically as a histogram and that the data appear ‘noisy’ in that they do not form a strong consensus at for an expected CNV value. A problem with the graphical representation of data arises when a large number of samples are to be explored or when the genome is subset into a large number of fractions, such as in windowing analyses. A numerical summary table provides the ratio of alleles observed in any genome or in any fraction of a genome. The problem of noisy data may in part be due to variants of low quality (i.e., technical error) or potential variation in ploidy throughout a genome or sub-genomic region (i.e., biological variation).

The challenge of identifying high quality variants and numerically summarizing them was addressed by our method of allele balance analysis (Knaus and Grünwald 2018). The data were quality filtered using the sequence depth of the most abundant allele for all variants in a genome. An 80% confidence interval was created to eliminate variants with the lowest 10% and highest 10% sequence coverage. This confidence interval was then applied to the second most abundant allele as well. The VCF file was further subset to only heterozygous positions. The allele balance ratio for each heterozygous variant was calculated by dividing the number of times the most abundant allele was sequenced by the number of times the most abundant allele and the second most abundant allele were sequenced, resulting in a proportion. Finally, 200,000 bp windows were made using the allele ratio data. This window size was chosen for *P. infestans* because it was sufficiently large to include a population of heterozygous positions (we observed a heterozygous position every 1-2 kbp) but small enough to obtain fine scale resolution. The data were then assigned to bins ranging from 0 to 1 that are of width 0.02 and the bin with the greatest density was used as a summary for the window. This is analogous to the modal frequency. This summary was then categorized to a ploidy level by assigning it to the closest expected ratio (i.e., 1/2, 2/3, 3/4, 4/5). Each genome was now summarized into windows of ploidy. In order to assign copy number to genes the coordinates of each gene were referenced in the windowed genome and the copy number of the window where the gene was located was used to assign a copy number to the gene. This is critical because we do not expect most genes to contain enough heterozygous positions to infer an accurate estimate of copy number. Once a copy number was determined a confidence in this estimate was made by subtracting the observed proportion from the determined proportion and dividing by the bin width so that the value ranges from zero to one. Calculations were performed in R (R Core Team 2018) and using vcfR (Knaus and Grünwald 2017, 2018).

### Gene loss based on breadth of coverage

In order to determine gene loss we measured breadth of coverage (BOC) for each gene in each genome. We used SAMtools mpileup (Li et al. 2009) to count per position sequence coverage over all 18,179 genes in the *P. infestans* T30-4 genome (Haas et al. 2009). From this data, the number of positions that were sequenced at least once and a median of coverage were collected. Breadth of coverage was calculated by dividing the number of positions that were sequenced at least once by the gene length (i.e., the proportion of positions sequenced in a gene). We used a BOC of zero to indicate the loss of a gene.

### Gene class and density

Published gene annotations (Haas et al. 2009) were used to assign genes to gene classes (core, pseudogene, RxLR, etc.). The flanking intergenic region (FIR) lengths (i.e., intergenic distances) were calculated using a previously available script (https://figshare.com/articles/Calculate_FIR_length_perl_script/707328). This information was used to create FIR plots for individuals and populations from Mexico, South America, and the lineage US-1 using R (R Core Team 2018) and ggplot2 (Wickham 2009). In order to explore whether genes of a particular class from populations from Mexico, South America, and the lineage US-1 were enriched for a particular copy number the genes were assigned a copy number (based on allele balance) and plotted as box and whisker plots using ggplot2 (Wickham 2009). In order to visualize whether genes determined to have three copies were more abundant in the gene dense or gene sparse portion of the genome FIR plots were created as above except using core orthologous genes that were determined to have three copies.

### Copy number variation in other species of *Phytophthora*

In order to address whether copy number variation occurred in other species of *Phytophthora* we queried NCBI for samples that had Illumina sequence data as well as an assembled genome reference for the species. These data were processed as the *P. infestans* data were processed. In order to visualize these data in a phylogenetic context a tree from Martin et al. (2014) was obtained from TreeBase (Vos et al. 2012). The data were then plotted in R (R Core Team 2018).

## Acknowledgements

Val Fieland, Karan Fairchild, Meg Larsen and Caroline Press provided much appreciated technical support. We much appreciate receiving data and cultures from Bill Fry and the USABlight community (https://usablight.org/) used for Figure supplement 1. We thank the Center for Genome Research and Biocomputing (CGRB) at Oregon State University for genome sequencing and support of our computational research on the CGRB cloud. This research was supported in part by US Department of Agriculture (USDA) Agricultural Research Service Grant 2072-22000-041-00-D and USDA National Institute of Food and Agriculture Grants 2011-68004-30154 and 2018-67013-27823. Mention of trade names or commercial products in this manuscript are solely for the purpose of providing specific information and do not imply recommendation or endorsement.

## 1 Dominant and ephemeral lineages

Emergent clonal lineages of *P. infestans* occur as “dominant” and “ephemeral” lineages. Ephemeral lineages may be only observed once or for one season. Dominant lineages are abundant over several seasons and their emergence may be declared an epidemic. For example, the dominant lineage US-1 was once thought to be the single clonal lineage present globally (Goodwin et al., 1994). During the 1990s dominant lineages US-8 and US-11, lineages that have resistance to the fungicide metalaxyl, displaced US-1 to become the dominant lineages (Fry and Goodwin, 1997a,b). In 2009 a new epidemic with new lineages occurred (Fry et al., 2013; Hu et al., 2012). The dominant lineage US-23 was observed on potato in 2009 but was not abundant (Figure supplement 1). Since 2009 it has steadily become more abundant and is now the most frequently observed lineage on potato or tomato. In contrast, the ephemeral lineage US-24 was abundant in 2010 and 2011 on potato but has almost completely disappeared in subsequent years. As similar pattern of dominant and ephemeral lineages has been reported from Europe (Cooke et al., 2012).

**Figure Supplement 1.**
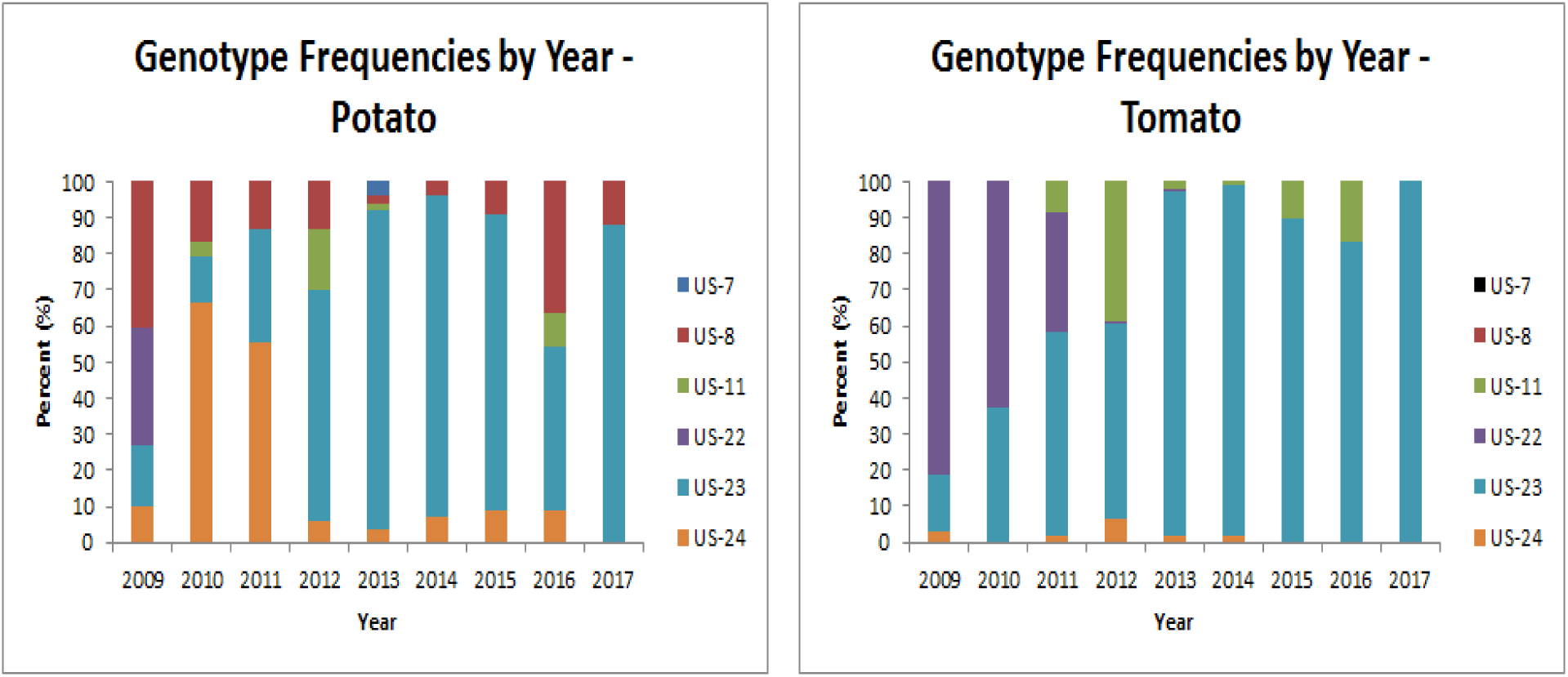
Lineages of *P. infestans* occur as dominant and ephemeral lineages. Stacked barcharts indicate the relative abundance of each lineage for each year. Data courtesy of William E. Fry (Cornell University) and USAblight (http://usablight.org/).

## 2 Read depth

The inference of copy number, including gene loss, is dependent on sequence depth. For example, the loss of a gene may be determined by a lack of sequence coverage. Similarly, the presence of three copies of a gene may be inferred when the sequence depth for alleles at a locus are present in multiples of 1/3. Insufficient sequence depth could result in the incorrect inference that a gene has been lost when it may actually be below the level of detection, perhaps because it is in a region of the genome that is difficult to sequence. Similarly, if a gene exists as two copies but is sequenced at 3X coverage it would only be possible to incorrectly infer that it has three copies. In order to address the issue of sufficient sequence depth we began by filtering the available genomes to only analyze genomes of relatively high sequence depth.

Sequenced read depth for genes was used to determine whether a sample had sufficient coverage for further analysis. Per position read depth was measured using SAMtools mpileup (Li et al., 2009). A GC corrected estimate of per gene sequence depth was then calculated following Raffaele et al. (2010):

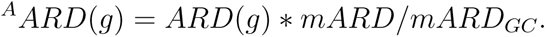

Here *ARD*(*g*) is the median read depth for a gene. Raffaele et al. (2010) reported average read depth but we have used median read depth to provide a robust estimate, but have retained the nomenclature of Raffaele and colleagues. *mARD* is the median read depth over all genes and *mARD_GC_* is the median read depth for genes in the same GC percentile as the gene being evaluated. This results in a GC adjusted average read depth *^A^ARD*(*g*).

A threshold of 12X coverage was used to determine suitability for analysis. Because our estimate of copy number relied on calculating the frequency of allele depth we chose a number that was greater than 10. Also, because we expected some samples to have three copies we chose a threshold that would divide evenly by two and three. Lastly, the choice of threshold was a compromise between attempting to select the highest coverage samples available, but also retaining as many samples as possible. The samples KM177497, M.0182897, and M.0182904 (Yoshida et al., 2013) were omitted because they did not appear to have enough data to process. Violin plots of were created of *^A^ARD*(*g*) for each available sample using ggplot2 (Wickham, 2009).

The samples T30-4, PIC99189, 90128, PIC99167, PIC99114, F18 (Raffaele et al., 2010), M.0182897, M.0182898, M.0182900, M.0182903, M.0182904, M.0182906, M.0182907, KM177497, KM177500, KM177509, KM177512, KM177513, KM177514, KM177517, KM177548, P11633, P12204, P1362, P6096 (Yoshida et al., 2013), Pi1845A, Pi1845B, Pi1876, Pi1882 (Martin et al., 2013), Kew123, P3681, P6629, P6635, PHU006, PCZ050, P10650, P3873, EC3394 (Martin et al., 2015), PIC97136, PIC97442 (this report) were considered but omitted based on the criterion that they had a genic AARD less than 12.

**Figure Supplement 2A.**
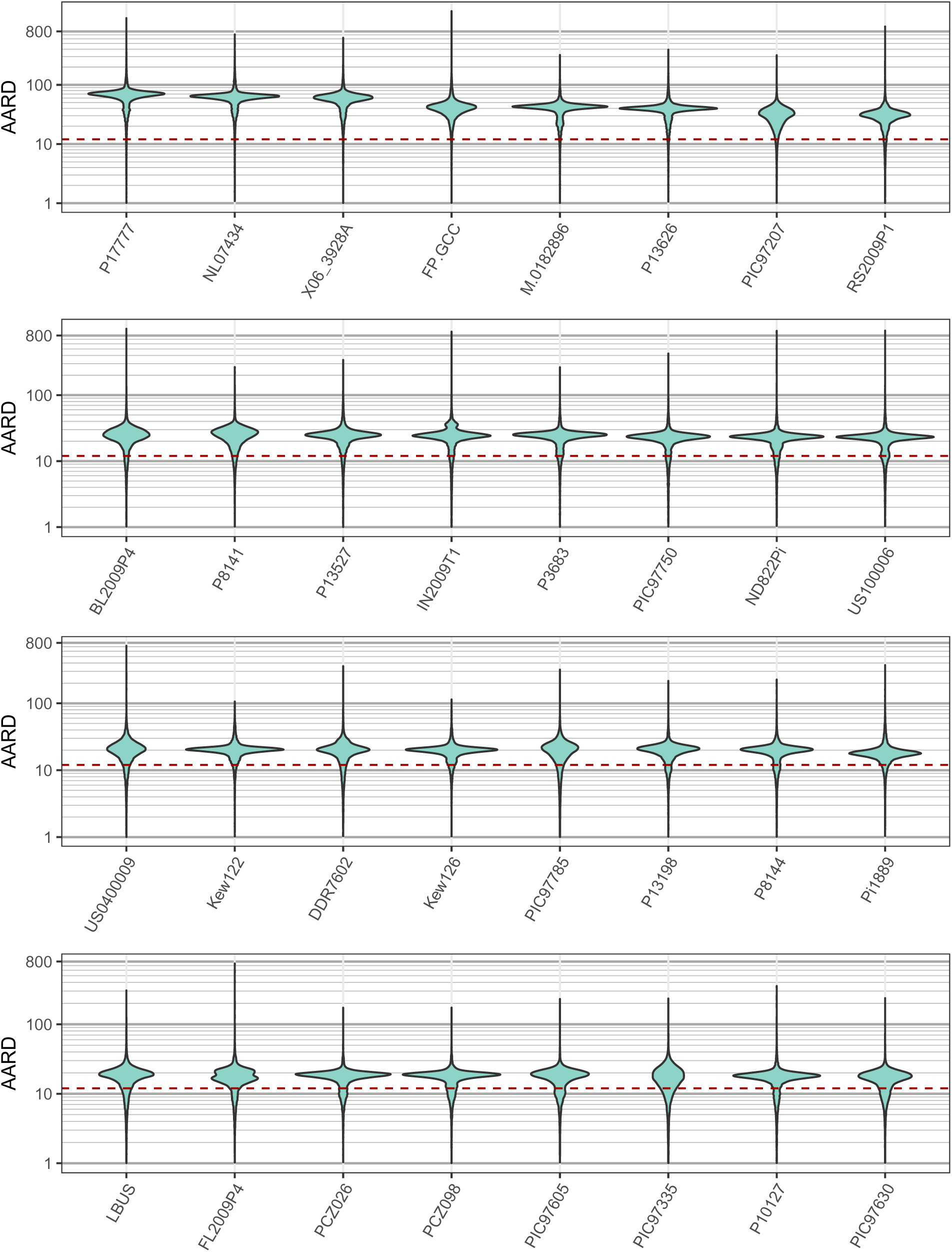
Violin plot of adjusted average read depth of for all *P. infestans* genes. The threshold for inclusion (12X) is indicated by a dashed red line.

**Figure Supplement 2B.**
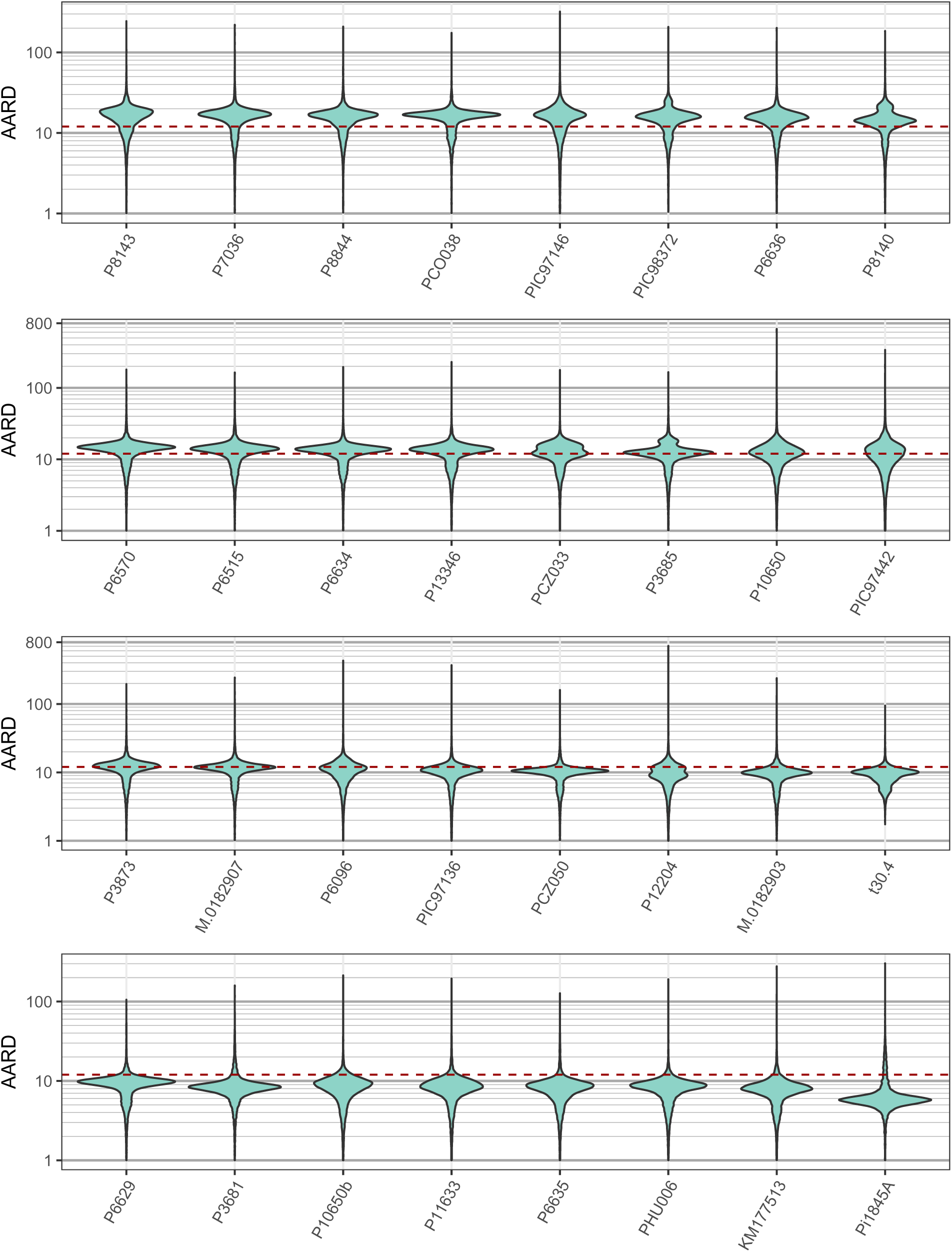
Violin plot of adjusted average read depth of for all *P. infestans* genes. The threshold for inclusion (12X) is indicated by a dashed red line.

**Figure Supplement 2C.**
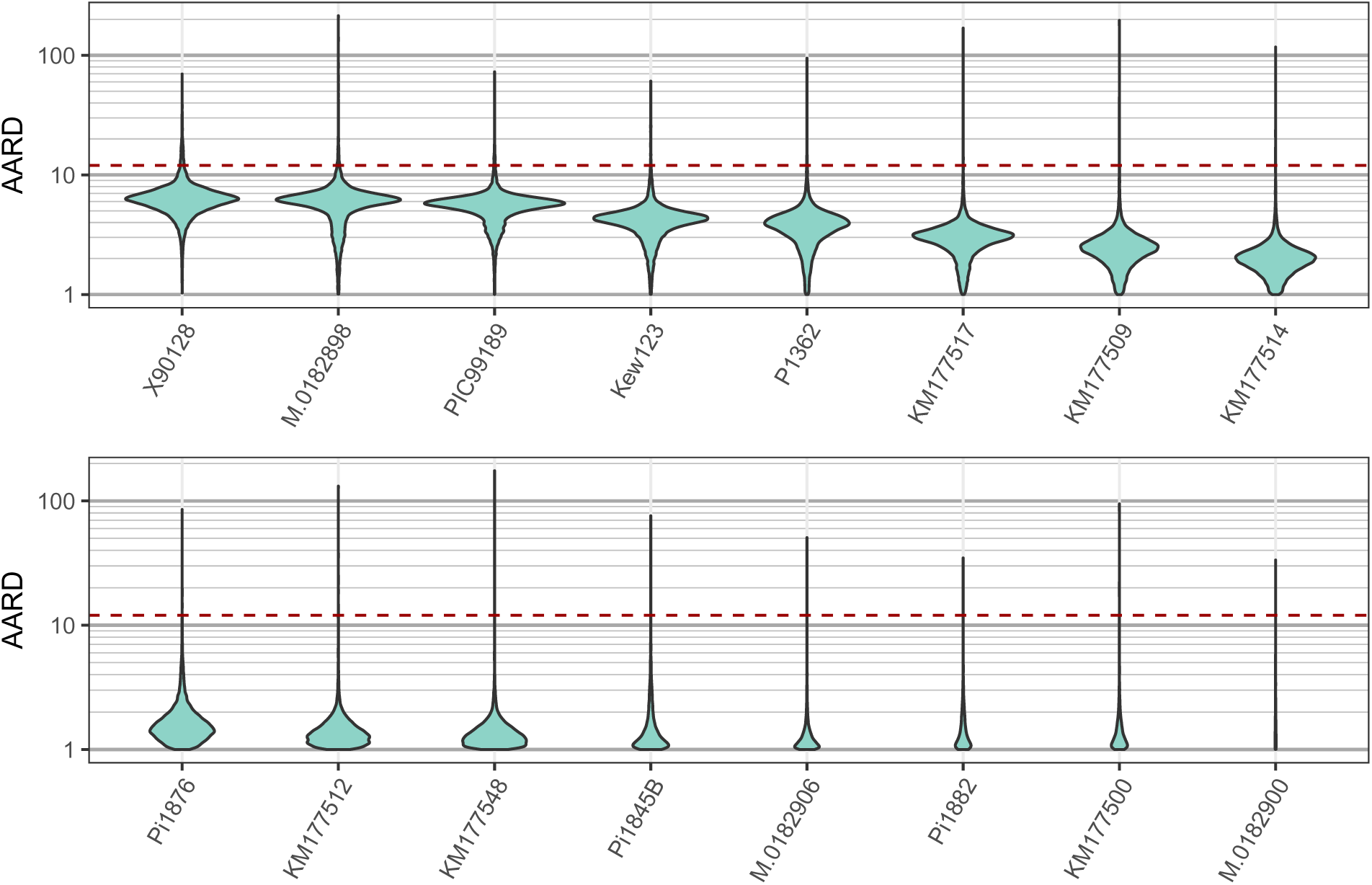
Violin plot of adjusted average read depth of for all *P. infestans* genes. The threshold for inclusion (12X) is indicated by a dashed red line.

## 3 *Phytophthora infestans* sample

**Table 1:** Isolates of *Phytophthora infestans* used in this report. Samples sequenced for this project are available at the sequence read archive (NCBI) as BioProject ID: PRJNA542680.

| Strain | Lineage | Year collected | Provenance | Mating type | SRA sample | Publication |
| --- | --- | --- | --- | --- | --- | --- |
| M-0182896 | HERB-1 | 1877 | Germany | NA | ERS241539 | Yoshida 2013 |
| Pi1889 | HERB-1 | 1889 | Germany | NA | ERS258007 | Martin 2013 |
| P8140 | HERB-1 | 1986 | Mexico, Toluca | A1 | SRS856518 | Martin 2016 |
| P13346 | HERB-1 | 2001 | Ecuador, Napo | A1 | SRS856501 | Martin 2016 |
| Kew126 | US-1 | 1952 | Britain | NA | SRS856492 | Martin 2016 |
| Kew122 | US-1 | 1955 | Britain | NA | SRS856493 | Martin 2016 |
| DDR7602 | US-1 | 1976 | Germany | A1 | ERS226848 | Yoshida 2013 |
| P8141 | US-1 | 1987 | Mexico | A1 | SRS856517 | Martin 2016 |
| P6515 | US-1 | 1989 | Peru | A1 | SRS856514 | Martin 2016 |
| LBUS5 | US-1 | 2005 | S. Africa | A1 | ERS226849 | Yoshida 2013 |
| RS2009P1 | US-8 | 2009 | USA, Pennsylvania | A2 | ERS258000 | Martin 2013 |
| US-0400009 | US-8 | NA | USA, New York | A2 | SAMN11619528 | This report |
| FP-GCC | US-11 | NA | USA, Florida | A1 | SAMN11635067 | This report |
| P10127 | US-18 | 2002 | USA, N.Carolina | A2 | ERS241587 | Yoshida 2013 |
| US-100006 | US-22 | NA | USA, Kentucky | A2 | SAMN11635068 | This report |
| P17777 | US-22 | 2009 | USA, New York | A2 | ERS226847 | Yoshida 2013 |
| IN2009T1 | US-22 | 2009 | USA, Pennsylvania | A2 | ERS258001 | Martin 2013 |
| BL2009P4 | US-23 | 2009 | USA, Pennsylvania | A1 | ERS258002 | Martin 2013 |
| FL2009P4 | US-23 | NA | USA, Florida | A1 | SAMN11635069 | This report |
| ND822Pi | US-24 | NA | USA, N. Dakota | A1 | SAMN11635070 | This report |
| P3685 | NA | 1983 | Mexico | A1 | SRS856523 | Martin 2016 |
| P6634 | NA | 1983 | Mexico | A1 | SRS856520 | Martin 2016 |
| P7036 | NA | 1983 | Mexico, Toluca | A2 | SRS856519 | Martin 2016 |
| P3683 | NA | 1983 | Mexico | A2 | SRS856524 | Martin 2016 |
| P8143 | NA | 1987 | Mexico, Toluca | A1 | SRS856515 | Martin 2016 |
| P6636 | NA | 1987 | Mexico, Toluca | A2 | SRS856499 | Martin 2016 |
| P8144 | NA | 1987 | Mexico, Toluca | A1 | SRS856516 | Martin 2016 |
| PIC97207 | NA | 1997 | Mexico | NA | SRS856506 | Martin 2016 |
| PIC97605 | NA | 1997 | Mexico | NA | SRS856505 | Martin 2016 |
| PIC97630 | NA | 1997 | Mexico | NA | SRS856504 | Martin 2016 |
| PIC97146 | NA | 1997 | Mexico | NA | SAMN11635071 | This report |
| PIC97335 | NA | 1997 | Mexico | A1 | SAMN11635072 | This report |
| PIC97785 | NA | 1997 | Mexico | NA | SAMN11635073 | This report |
| PIC97750 | NA | 1997 | Mexico | A2 | SAMN11635074 | This report |
| P10650 | NA | 1998 | Mexico, Toluca | A1 | SRS856503 | Martin 2016 |
| PIC98372 | NA | 1998 | Mexico | NA | SRS856498 | Martin 2016 |
| P8844 | NA | 1982 | Peru | A1 | SRS856526 | Martin 2016 |
| PCZ026 | PE6 | 1997 | Peru, Cusco | NA | SRS856510 | Martin 2016 |
| PCZ098 | EC1.3 | 1997 | Peru, Cusco | NA | SRS856507 | Martin 2016 |
| PCO038 | EC1 | 1997 | Peru | NA | SRS856512 | Martin 2016 |
| PCZ033 | EC1.2 | 1997 | Peru, Cusco | NA | SRS856509 | Martin 2016 |
| P13198 | EC1 | 1998 | Ecuador, Napo | A1 | SRS856502 | Martin 2016 |
| P13527 | NA | 2002 | Ecuador | A1 | ERS226844 | Yoshida 2013 |
| P13626 | NA | 2003 | Ecuador | A1 | ERS226845 | Yoshida 2013 |
| P6570 | NA | 1989 | The Netherlands | A1 | SRS856513 | Martin 2016 |
| 06_3928A | Blue13 | 2006 | England | A2 | ERS226850 | Cooke 2012 |
| NL07434 | NA | 2007 | The Netherlands | NA | ERS226846 | Yoshida 2013 |

## 4 Non-*P. infestans* sample

**Table 2:** Isolates of *Phytophthora* (*P. infestans* excluded) used in this report. Samples sequenced for this project are available at the sequence read archive (NCBI) as BioProject ID: PRJNA542680.

| Strain | Taxon | Year collected | Provenance | Mating type | SRA sample | Publication |
| --- | --- | --- | --- | --- | --- | --- |
| P13803 | *P. andina* | 2004 | Ecuador, Pichincha | A1 | SRS856497 | Martin 2016 |
| PaX | *P. andina* | NA | NA | NA | SRS856496 | Martin 2016 |
| Ipom_2.4 | *P. ipomoeae* | 1999 | Mexico | NA | SAMN11635080 | This report |
| Pic99167 | *P. ipomoeae* | 1999 | Mexico | NA | SAMN11635081 | This report |
| 00Ip05 | *P. ipomoeae* | 2000 | Mexico | NA | SAMN11635082 | This report |
| PI-07-097 | *P. ipomoeae* | 2004 | Mexico, Michoacan | NA | SAMN11635083 | This report |
| Ipom_1.2 | *P. ipomoeae* | 1999 | Mexico | NA | SAMN11635084 | This report |
| P7722 | *P. mirabilis* | 1992 | USA, California | A1 | ERS241588 | Yoshida 2013 |
| CBS.678.85 | *P. mirabilis* | NA | Mexico | NA | SAMN11635075 | This report |
| P3001 | *P. mirabilis* | 1998 | Mexico | NA | SAMN11635076 | This report |
| PIC99114b | *P. mirabilis* | 1998 | Mexico | NA | SAMN11635077 | This report |
| 00M 410 | *P. mirabilis* | 2000 | Mexico, D.F. | NA | SAMN11635078 | This report |
| PI-07-100 | *P. mirabilis* | 2003 | Mexico, Michoacan | NA | SAMN11635079 | This report |
| INRA-310 | *P. parasitica* | NA | NA | NA | SRS348810 | The Broad |
| J01A1 | *P. parasitica* | NA | NA | NA | NA | The Broad |
| P10297 | *P. parasitica* | NA | USA, Florida | A2 | SRS473780 | The Broad |
| P1569 | *P. parasitica* | NA | USA, California | A1 | SRS473781 | The Broad |
| P1976 | *P. parasitica* | 1976 | USA, California | A2 | SRS473782 | The Broad |
| Pc1E8 | *P. capsici* | 1987 | Taiwan, Taipei | NA | SRS2114629 | NA |
| Pc33e | *P. capsici* | 1995 | Taiwan, Tainan | NA | SRS2114628 | NA |
| Pc337 | *P. capsici* | 2016 | Taiwan, Yunlin | NA | SRS2115143 | NA |
| Pc389 | *P. capsici* | 2016 | Taiwan, Hualien | NA | SRS2115103 | NA |

## 5 Allele balance: *P. infestans*

Data were processed as in the section ‘Gene copy number inference based on allele balance’ and in Knaus and Grünwald (2018). Allele balance, the frequency at which each allele was sequenced, was calculated for each available sample. A genomic VCF file (g.VCF) was created for each sample using GATK (McKenna et al., 2010; DePristo et al., 2011). Each g.VCF file was read into R (R Core Team, 2017) for processing with vcfR (Knaus and Grünwald, 2017). The allele depth (AD) and genotypes were extracted from the vcfR object and the first and second most abundant alleles were extracted from the allele depth. The genotype information was used to subset the allele depth information to only heterozygous positions. The 15th and 85th percentiles were calculated for each sample and each alelle (the first and second most abundant alleles) and used as an inclusion threshold for depth filtering. A frequency for each allele was then calculated by dividing its allele depth by the sum of the first and second most abundant alleles. This information was used to plot each histogram. Samples are presented in the same order as in Figure Supplement 1A-C.

**Figure Supplement 3A.**
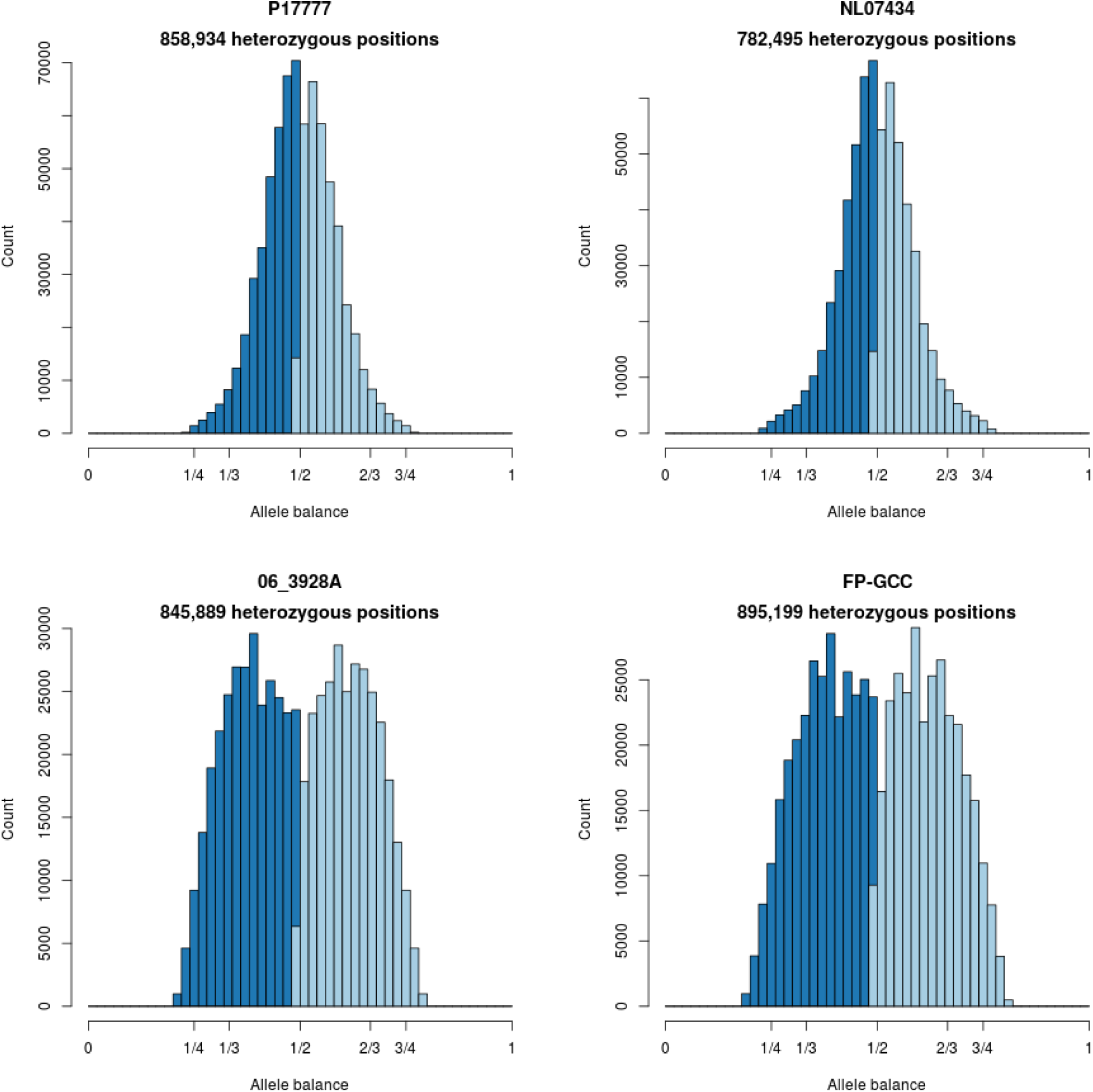
Histograms of allele balance for *P. infestans*.

**Figure Supplement 3B.**
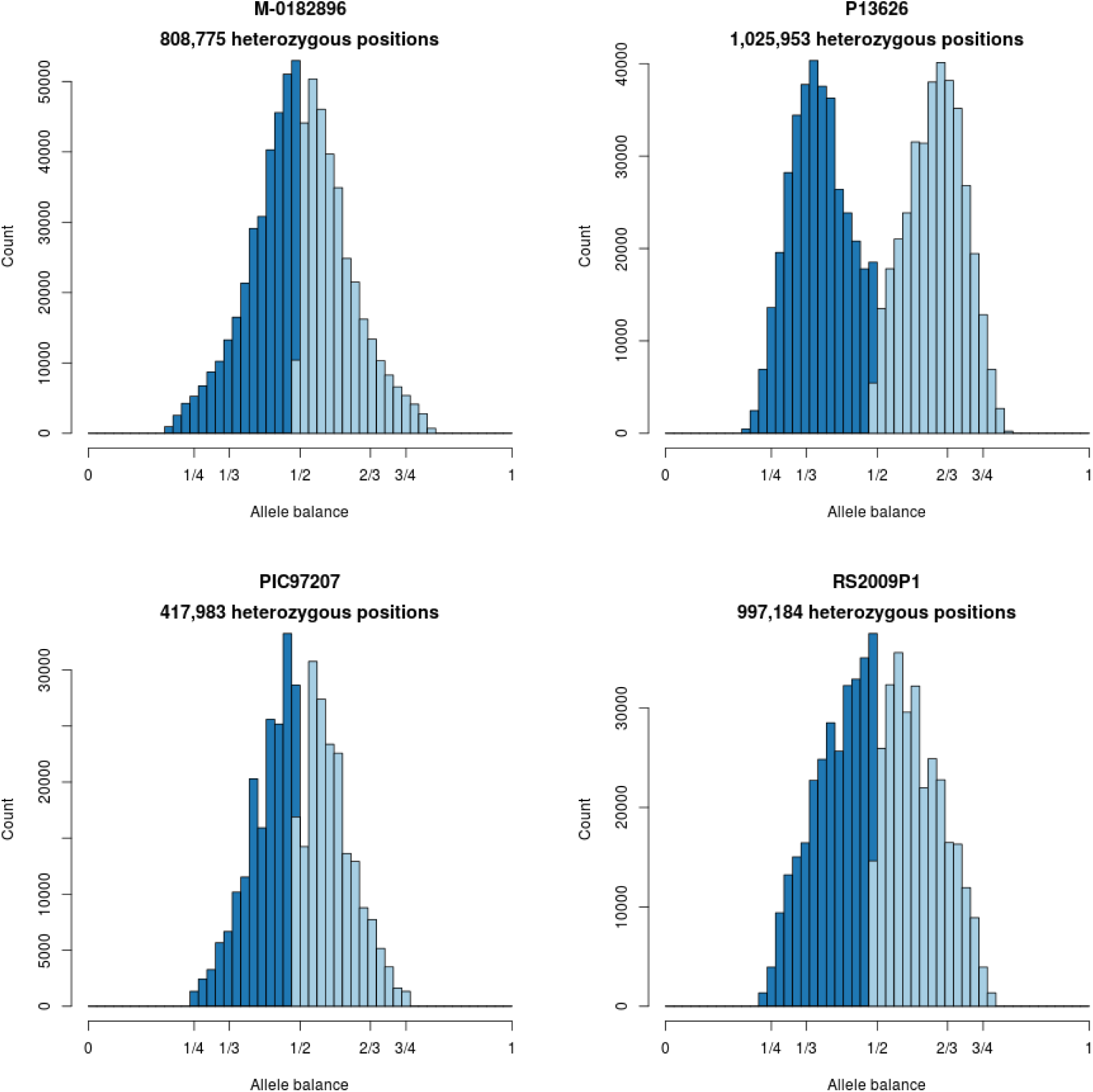
Histograms of allele balance for *P. infestans*.

**Figure Supplement 3C.**
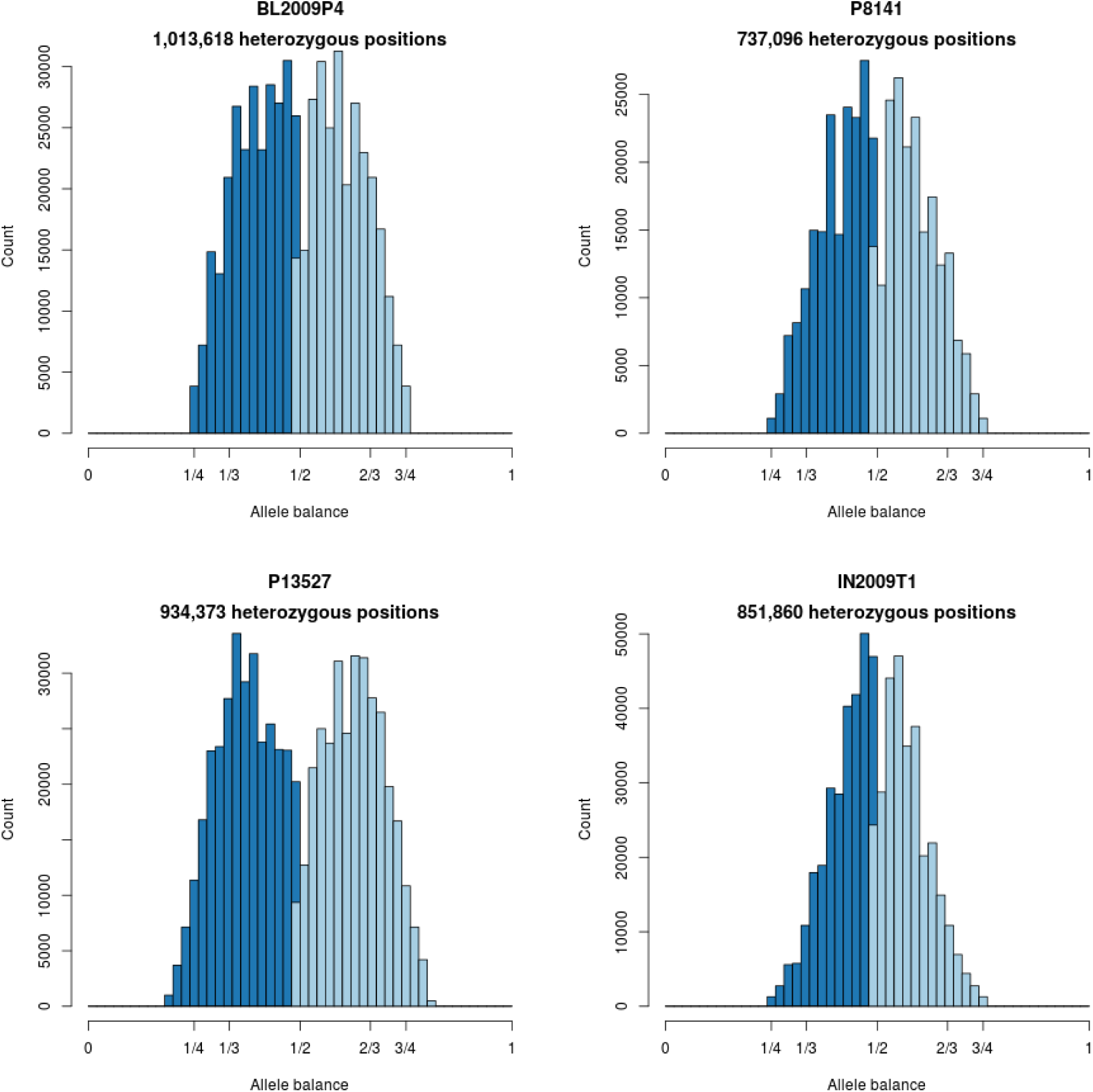
Histograms of allele balance for *P. infestans*.

**Figure Supplement 3D.**
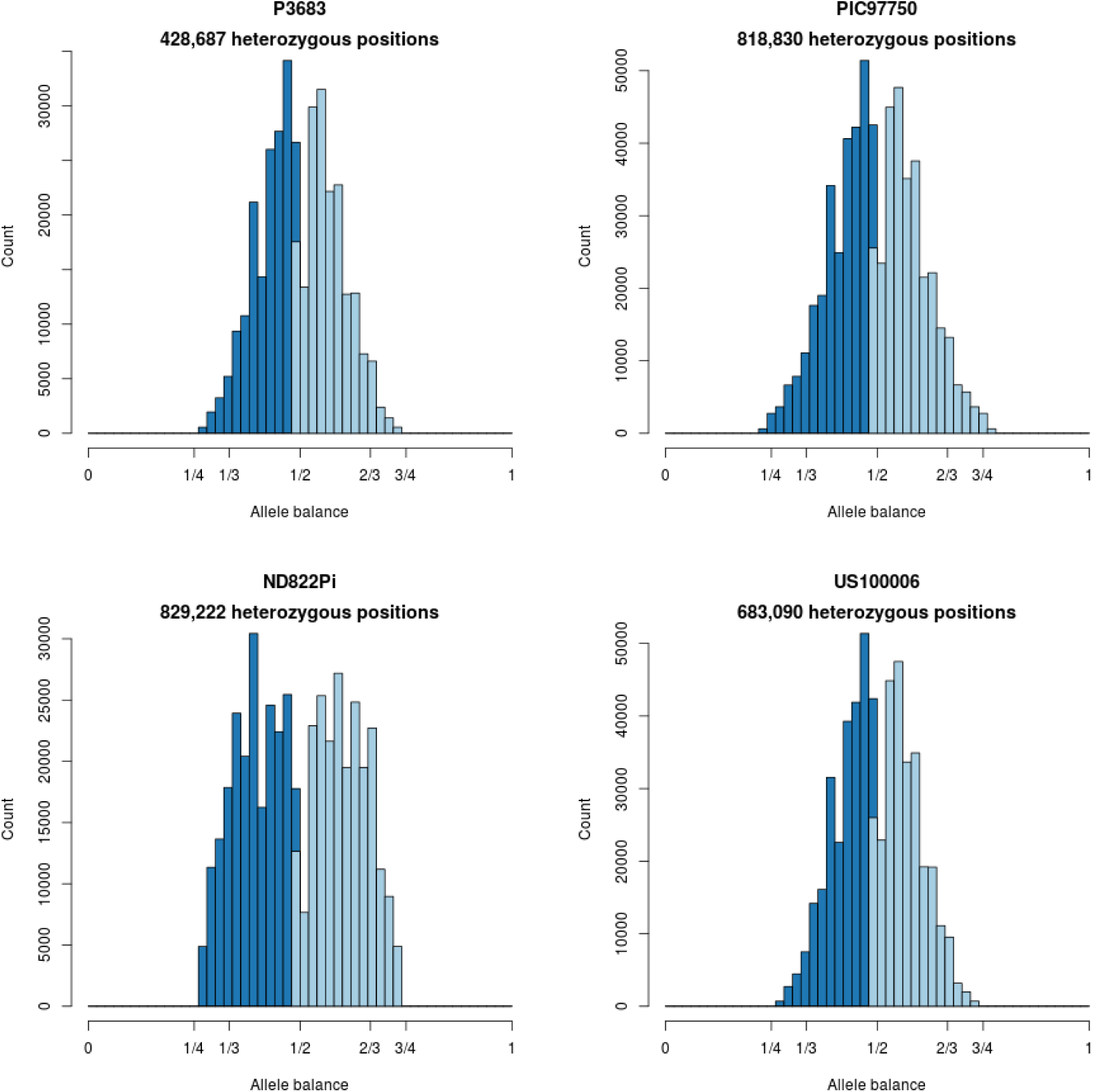
Histograms of allele balance for *P. infestans*.

**Figure Supplement 3E.**
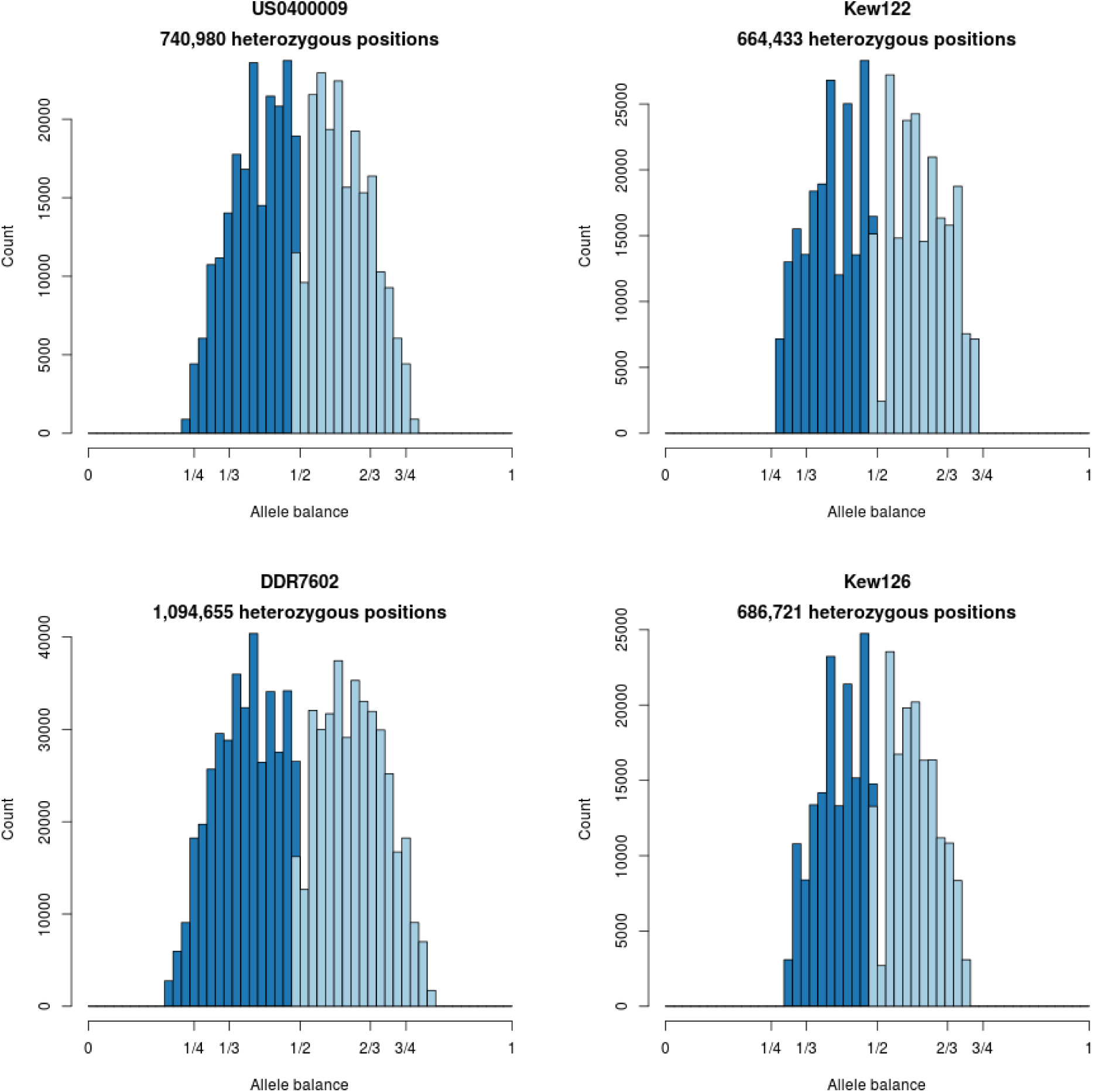
Histograms of allele balance for *P. infestans*.

**Figure Supplement 3F.**
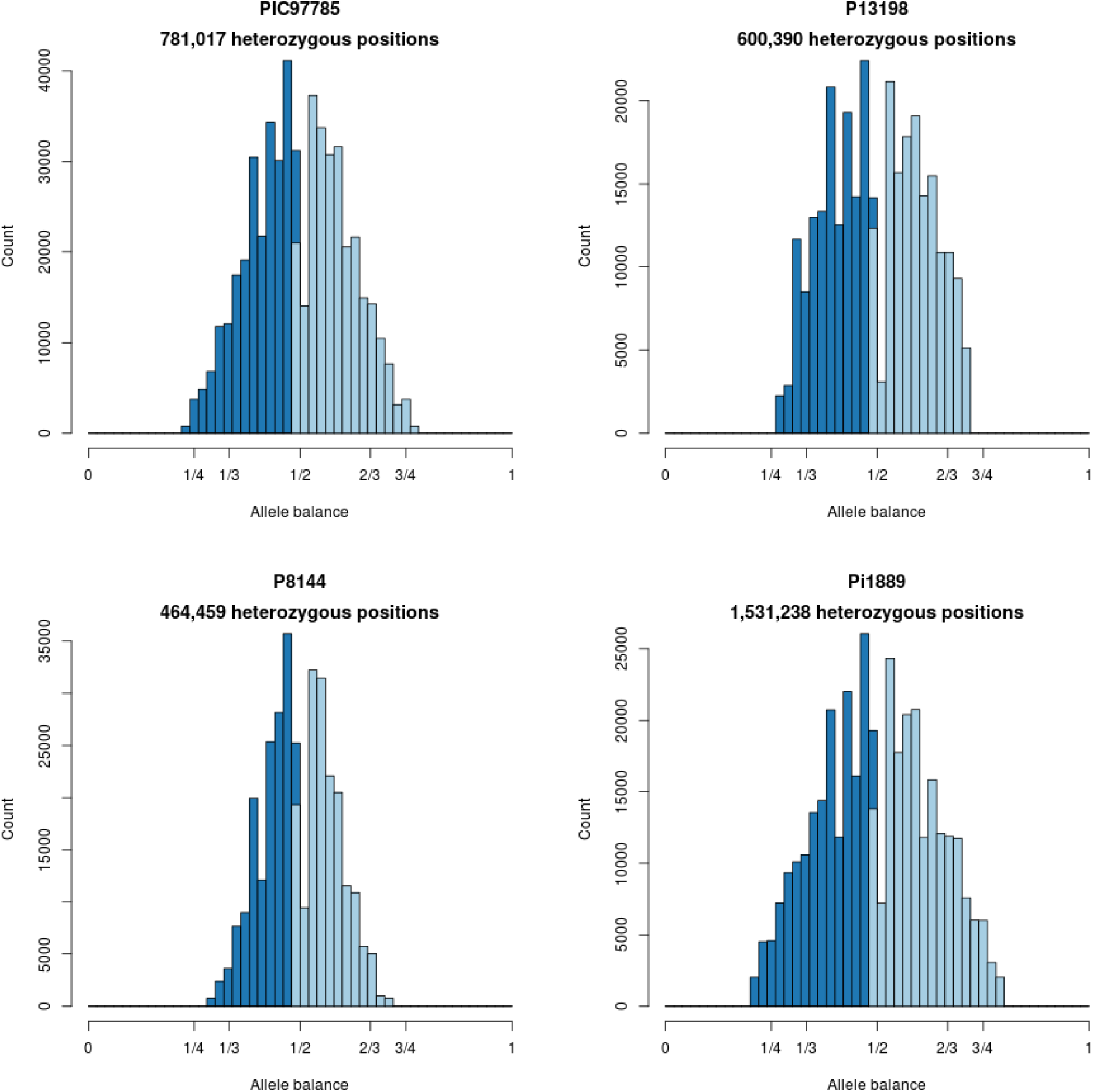
Histograms of allele balance for *P. infestans*.

**Figure Supplement 3G.**
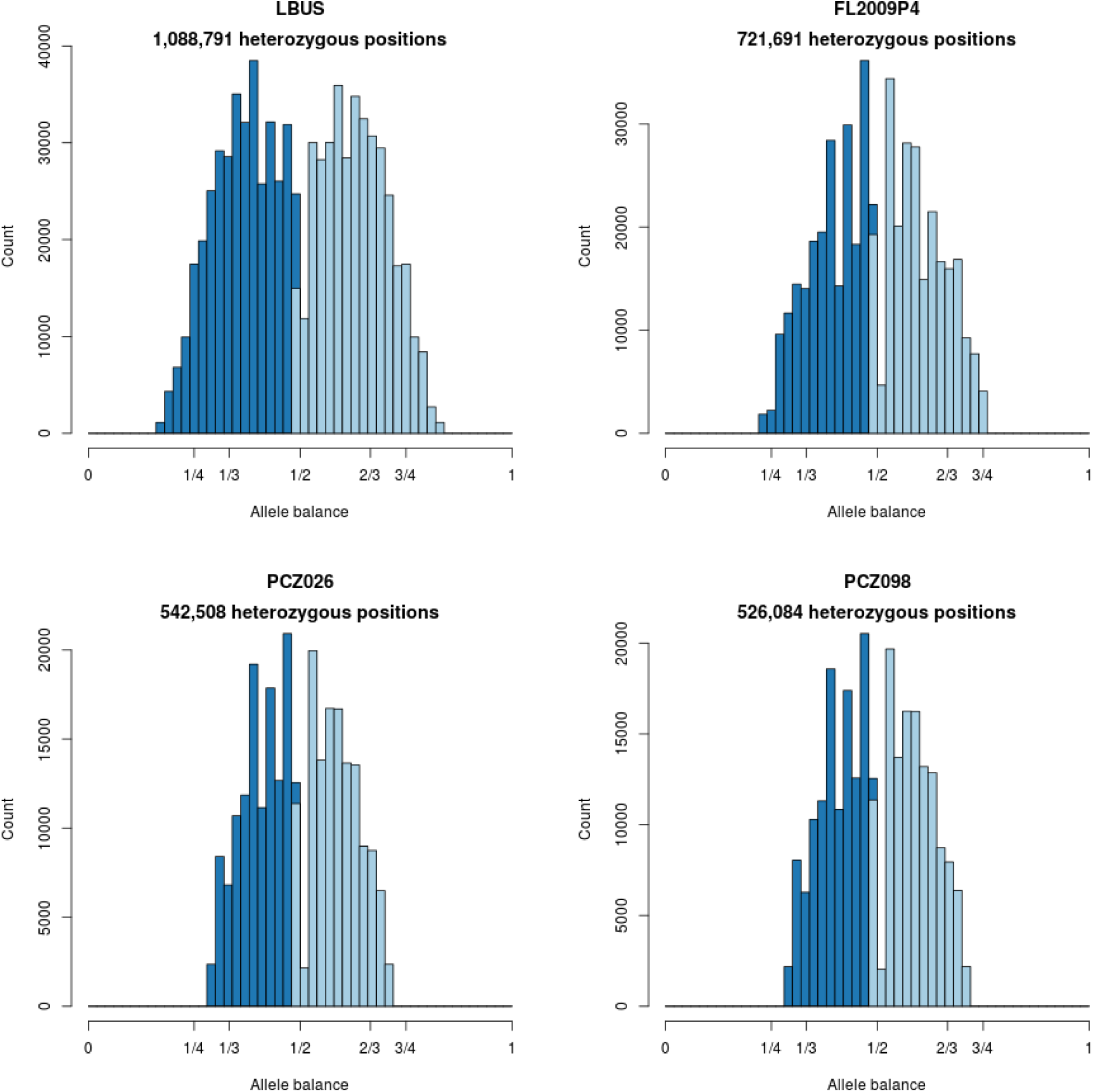
Histograms of allele balance for *P. infestans*.

**Figure Supplement 3H.**
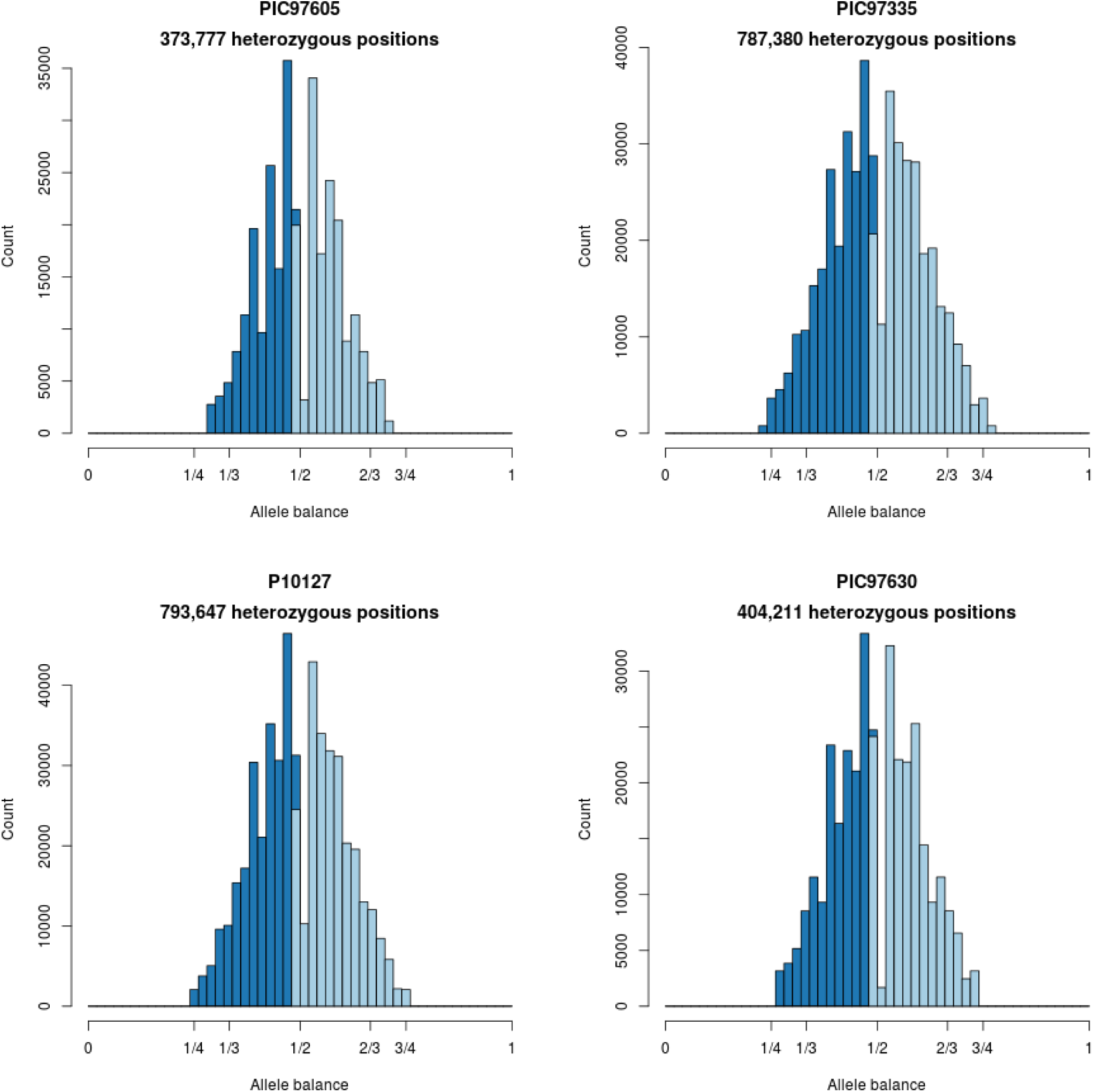
Histograms of allele balance for *P. infestans*.

**Figure Supplement 3I.**
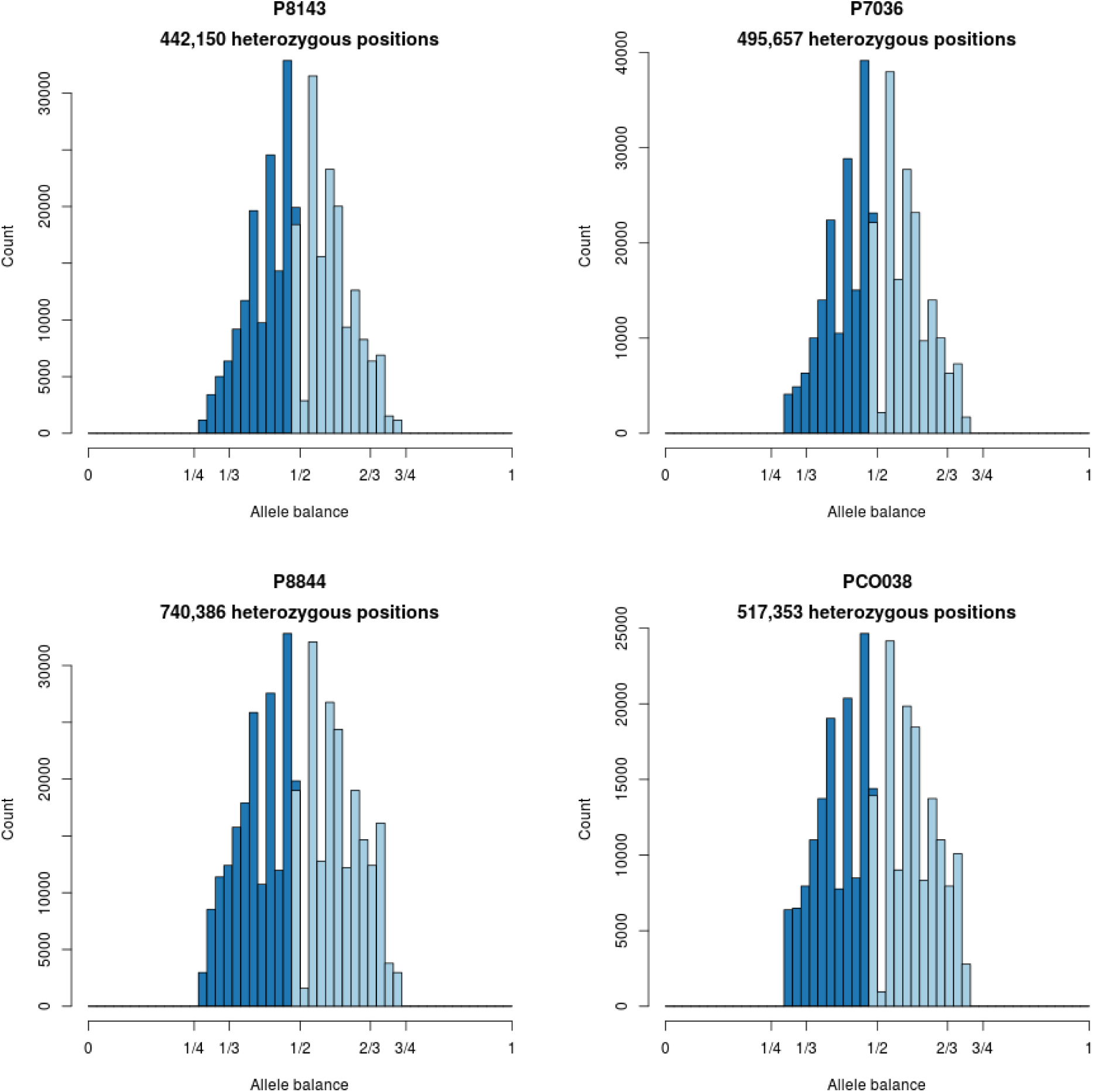
Histograms of allele balance for *P. infestans*.

**Figure Supplement 3J.**
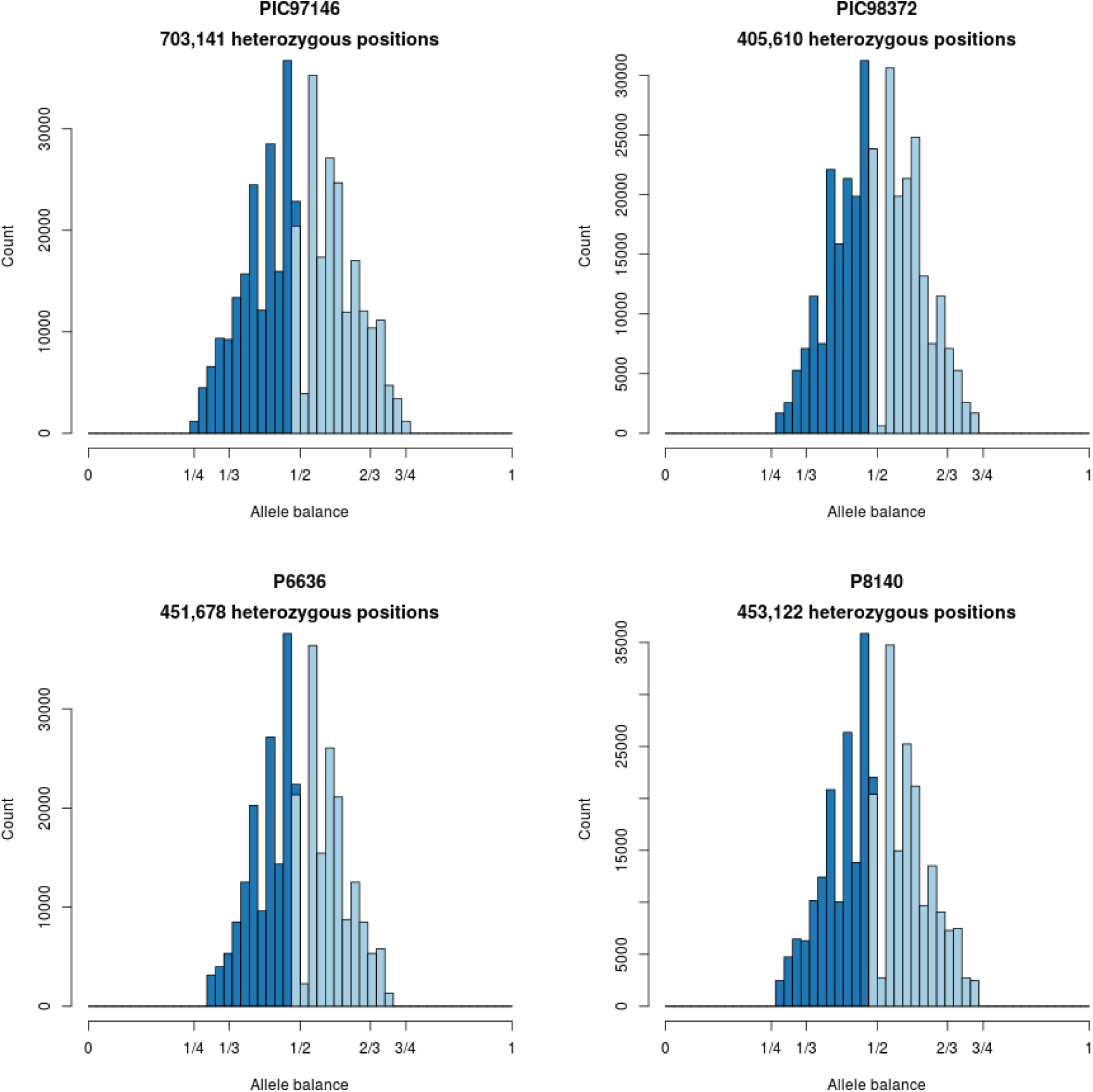
Histograms of allele balance for *P. infestans*.

**Figure Supplement 3K.**
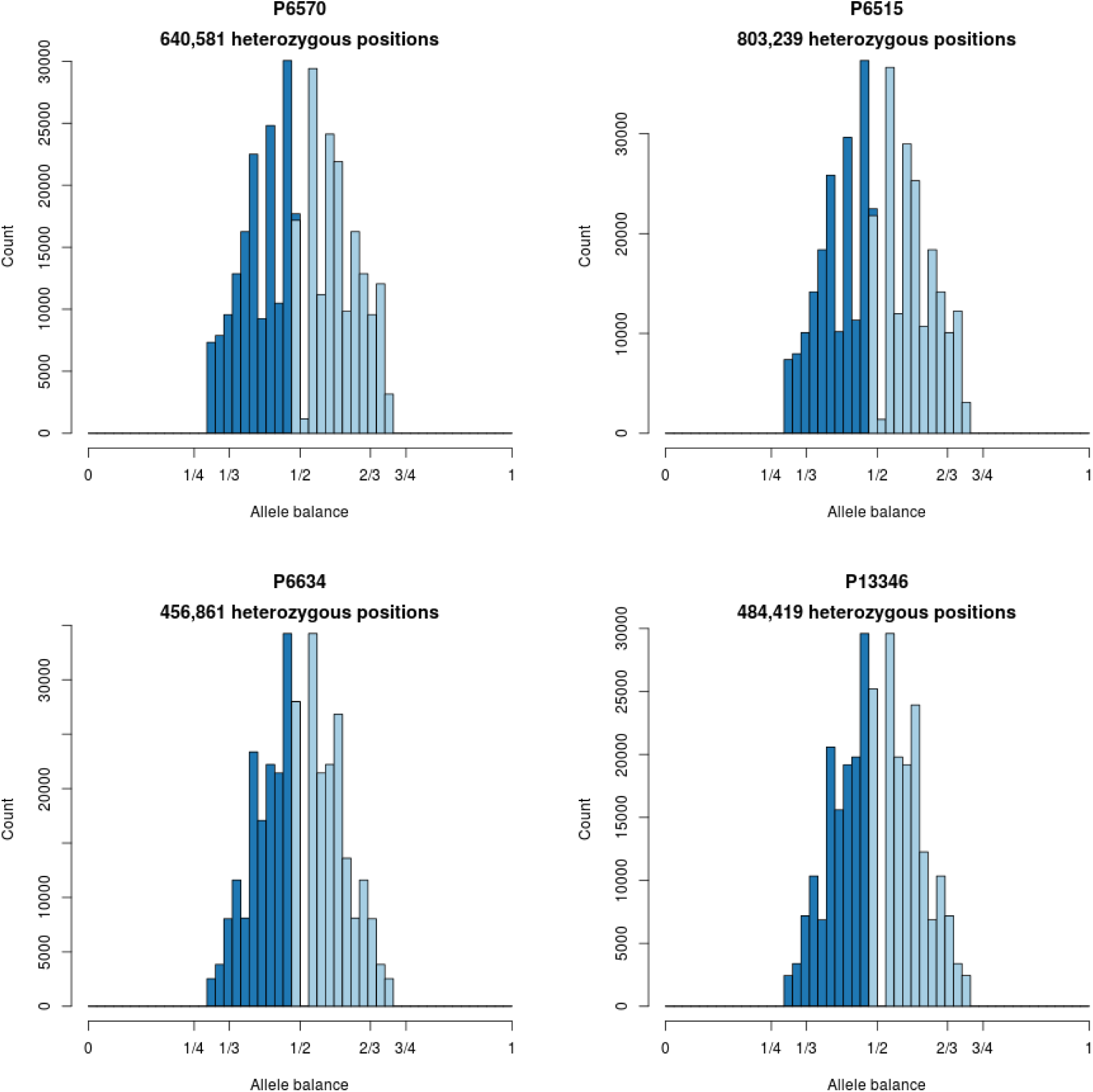
Histograms of allele balance for *P. infestans*.

**Figure Supplement 3L.**
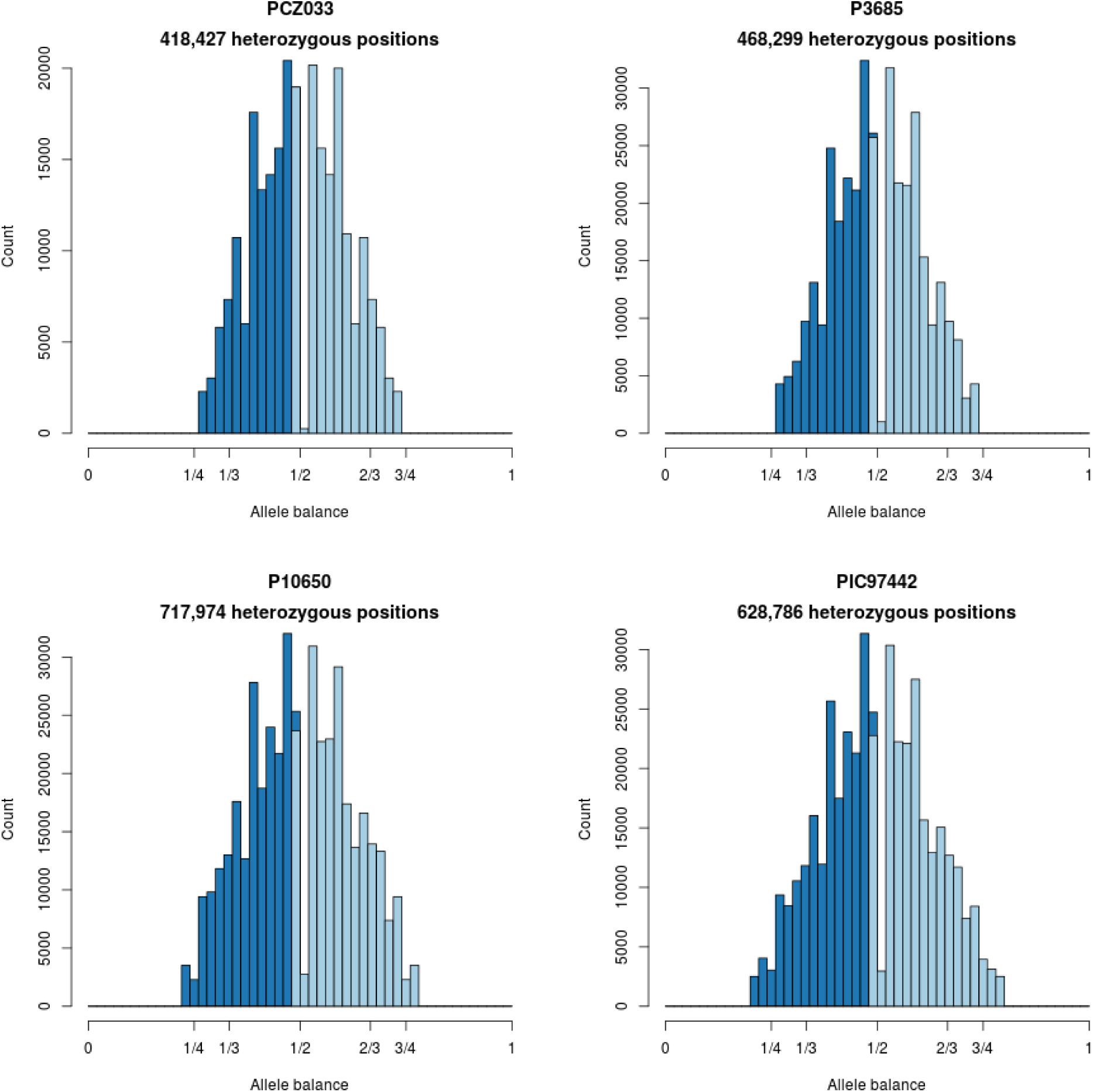
Histograms of allele balance for *P. infestans*.

**Figure Supplement 3M.**
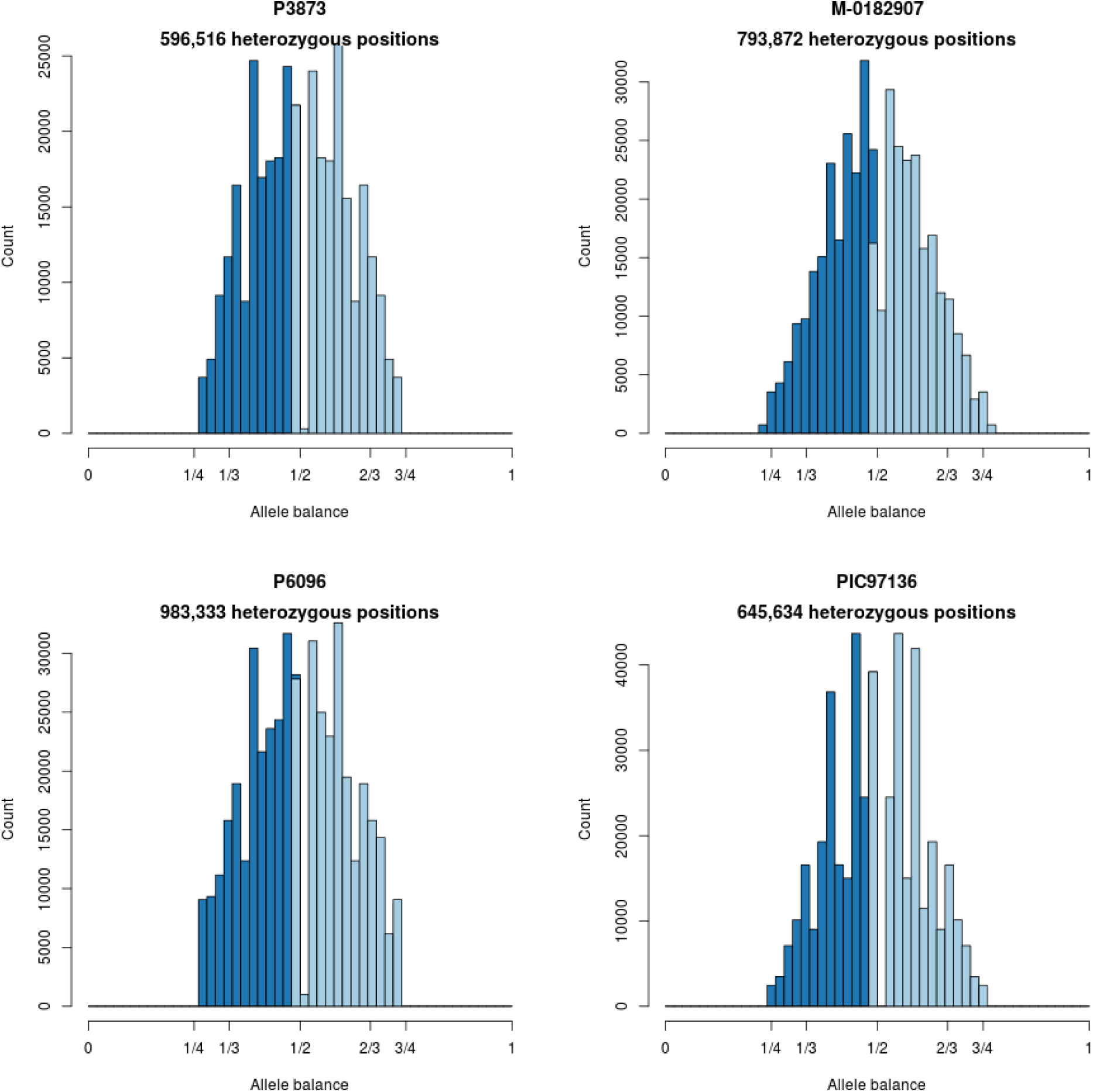
Histograms of allele balance for *P. infestans*.

**Figure Supplement 3N.**
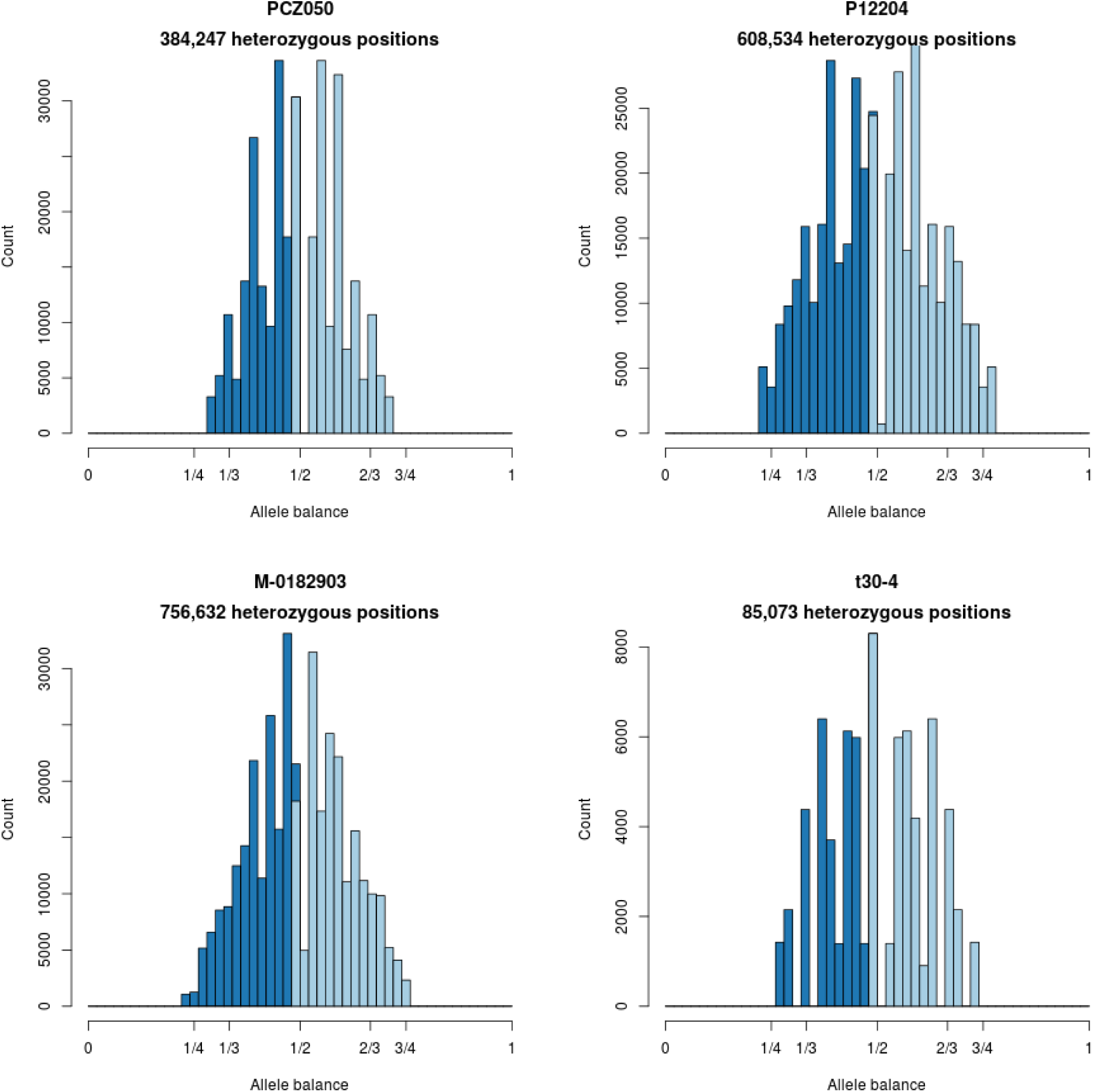
Histograms of allele balance for *P. infestans*.

**Figure Supplement 3O.**
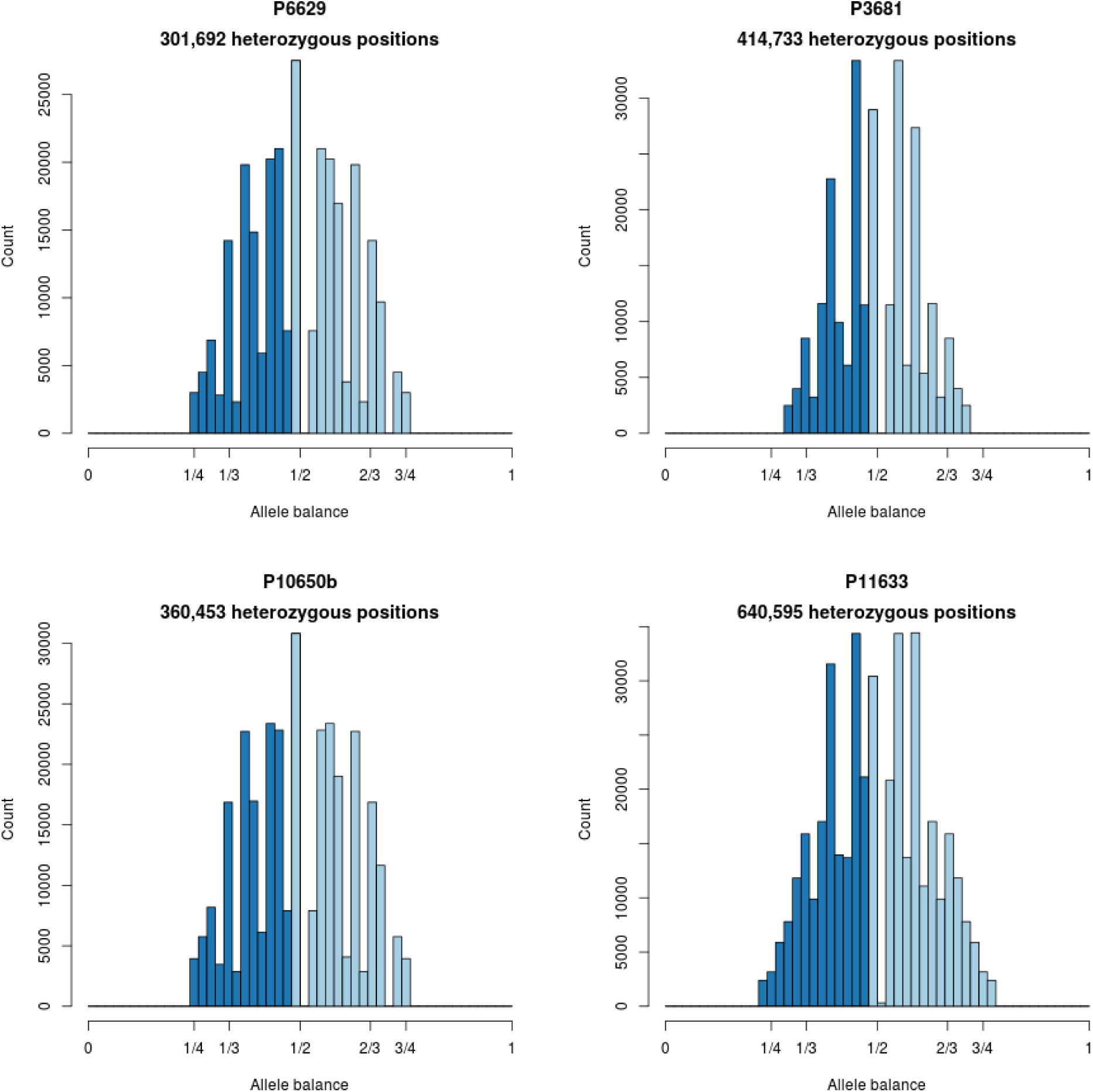
Histograms of allele balance for *P. infestans*.

**Figure Supplement 3P.**
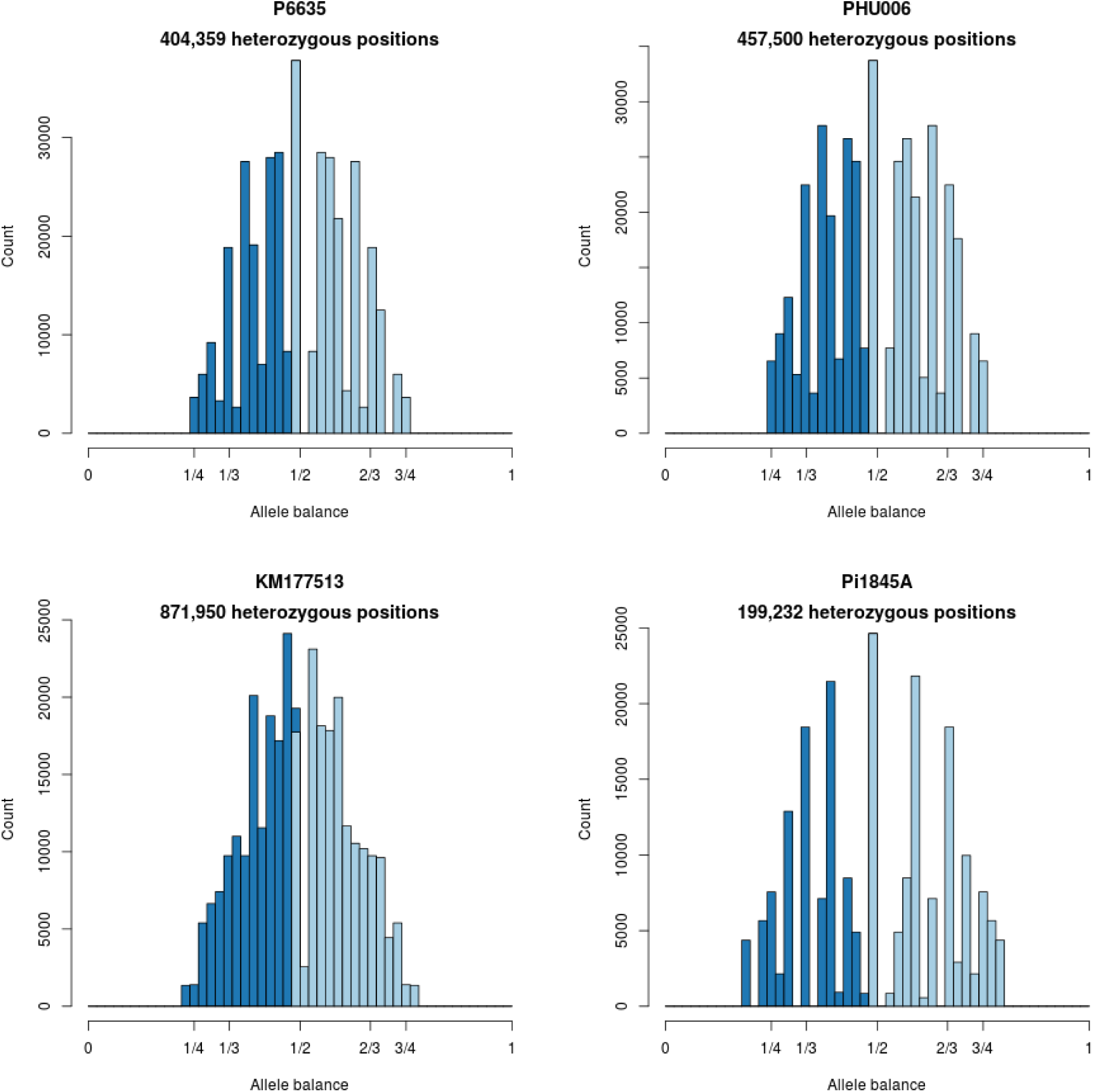
Histograms of allele balance for *P. infestans*.

**Figure Supplement 3Q.**
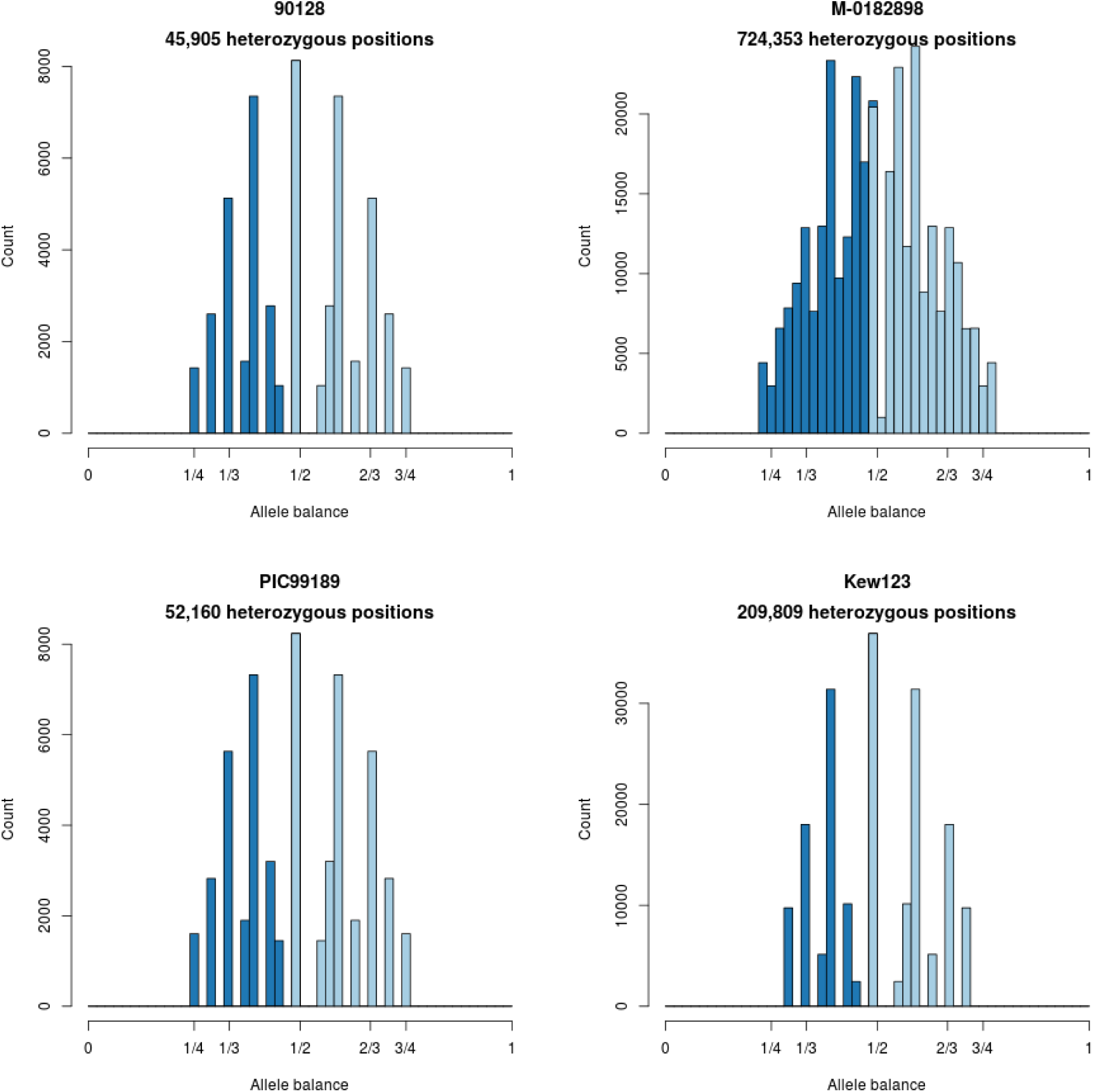
Histograms of allele balance for *P. infestans*.

**Figure Supplement 3R.**
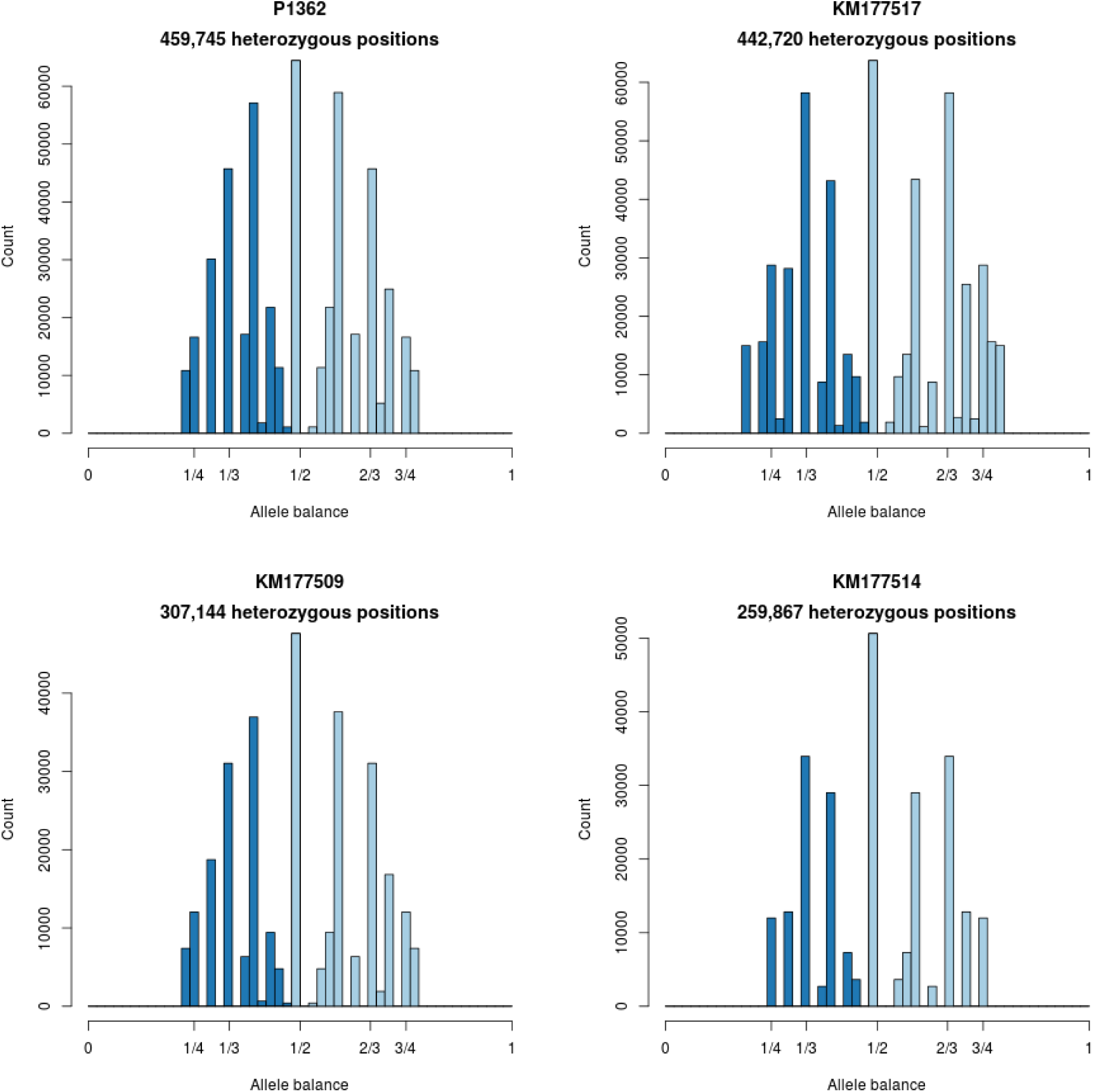
Histograms of allele balance for *P. infestans*.

**Figure Supplement 3S.**
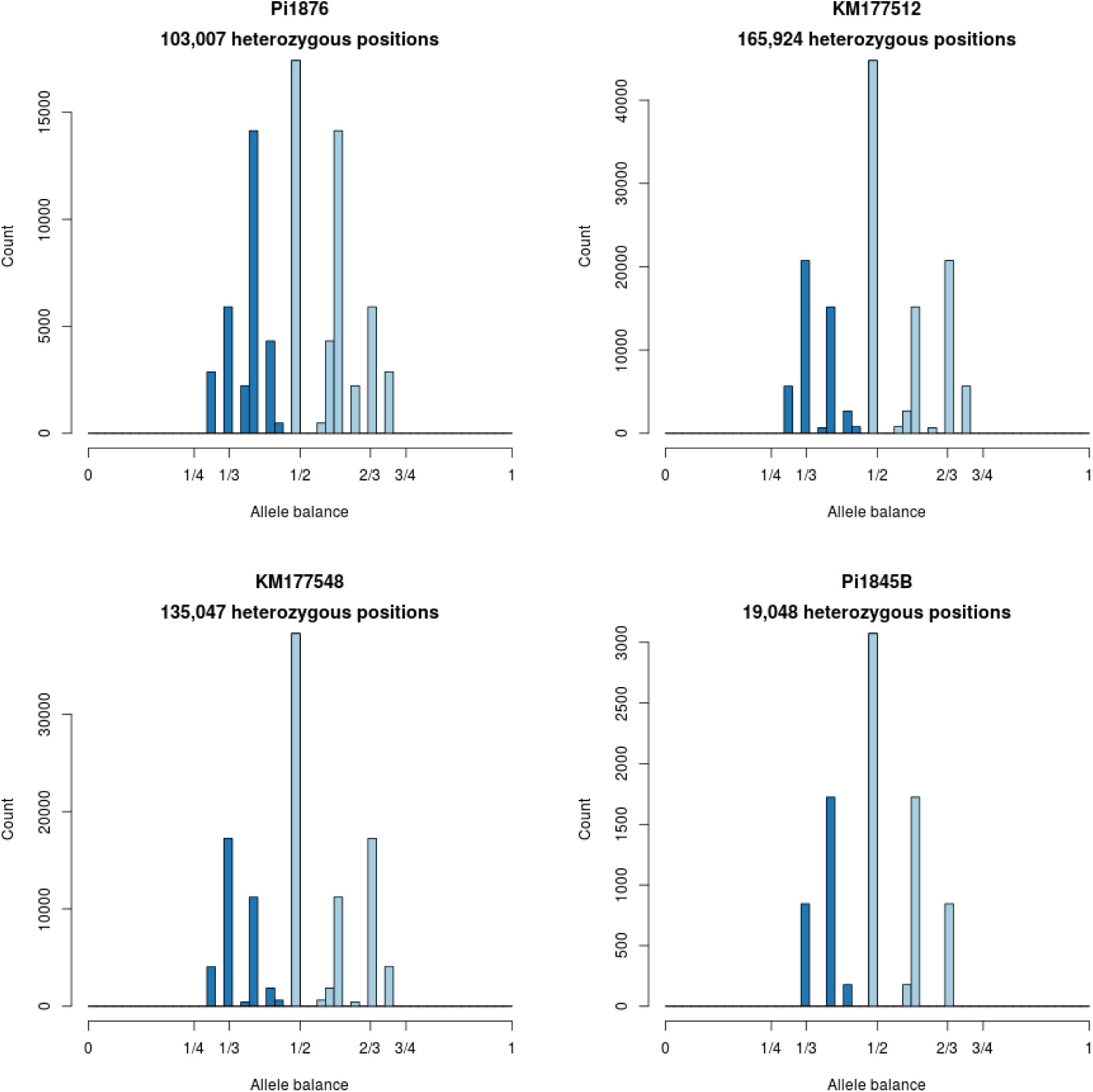
Histograms of allele balance for *P. infestans*.

**Figure Supplement 3T.**
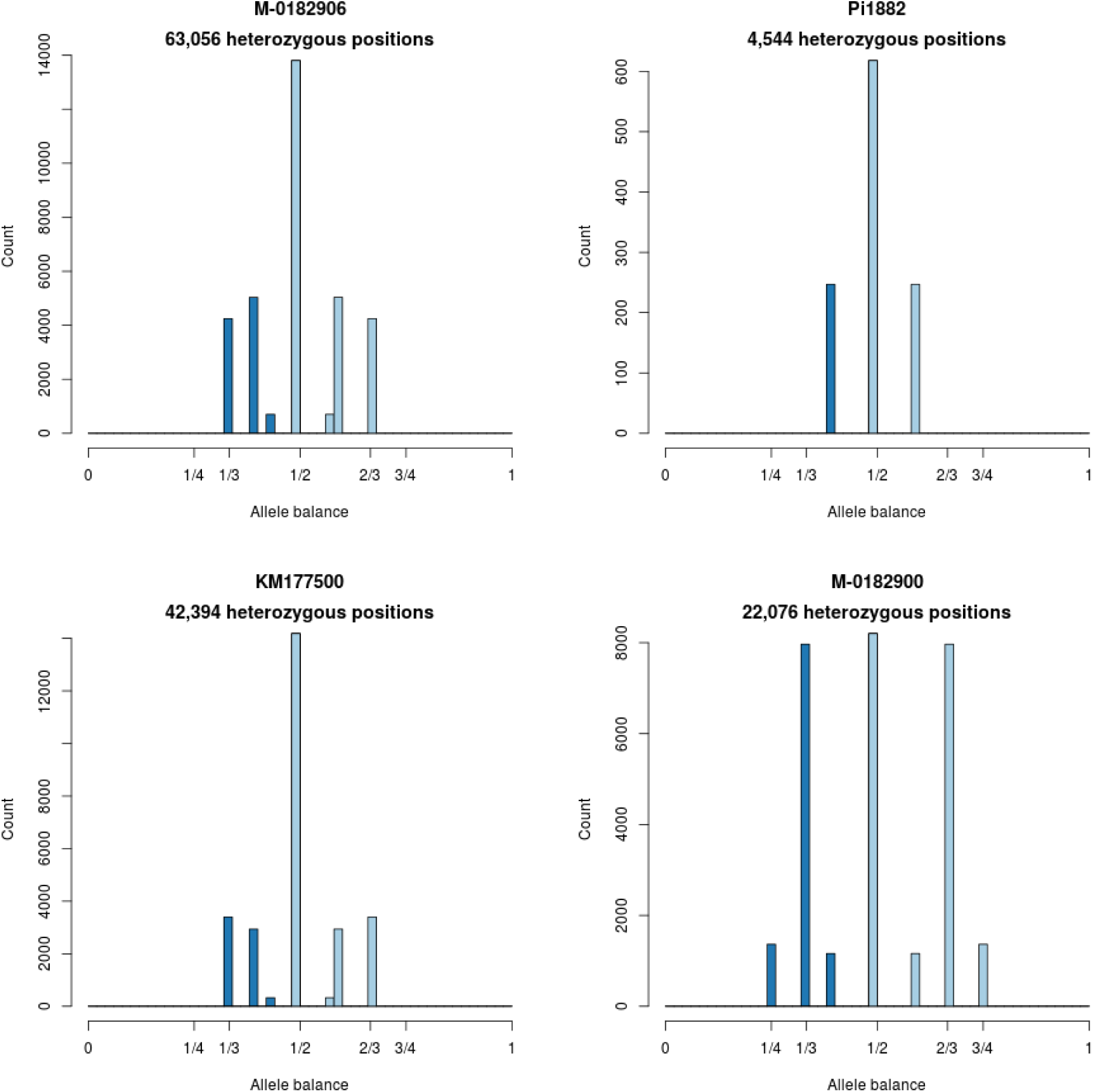
Histograms of allele balance for *P. infestans*.

## 6 Allele balance: non-*P. infestans* samples

Allele balance, the frequency at which each allele was sequenced, was calculated for each available sample. A genomic VCF file (g.VCF) was created for each sample using GATK (McKenna et al., 2010; DePristo et al., 2011). Each g.VCF file was read into R (R Core Team, 2017) for processing with vcfR (Knaus and Grünwald, 2017). The allele depth (AD) and genotypes were extracted from the vcfR object and the first and second most abundant alleles were extracted from the allele depth. The genotype information was used to subset the allele depth information to only heterozygous positions. The 15th and 85th percentiles were calculated for each sample and each alelle (the first and second most abundant alleles) and used as an inclusion threshold for depth filtering. A frequency for each allele was then calculated by dividing its allele depth by the sum of the first and second most abundant alleles. This information was used to plot each histogram. These methods are also described in Knaus and Grünwald (2018).

**Figure Supplement 4A.**
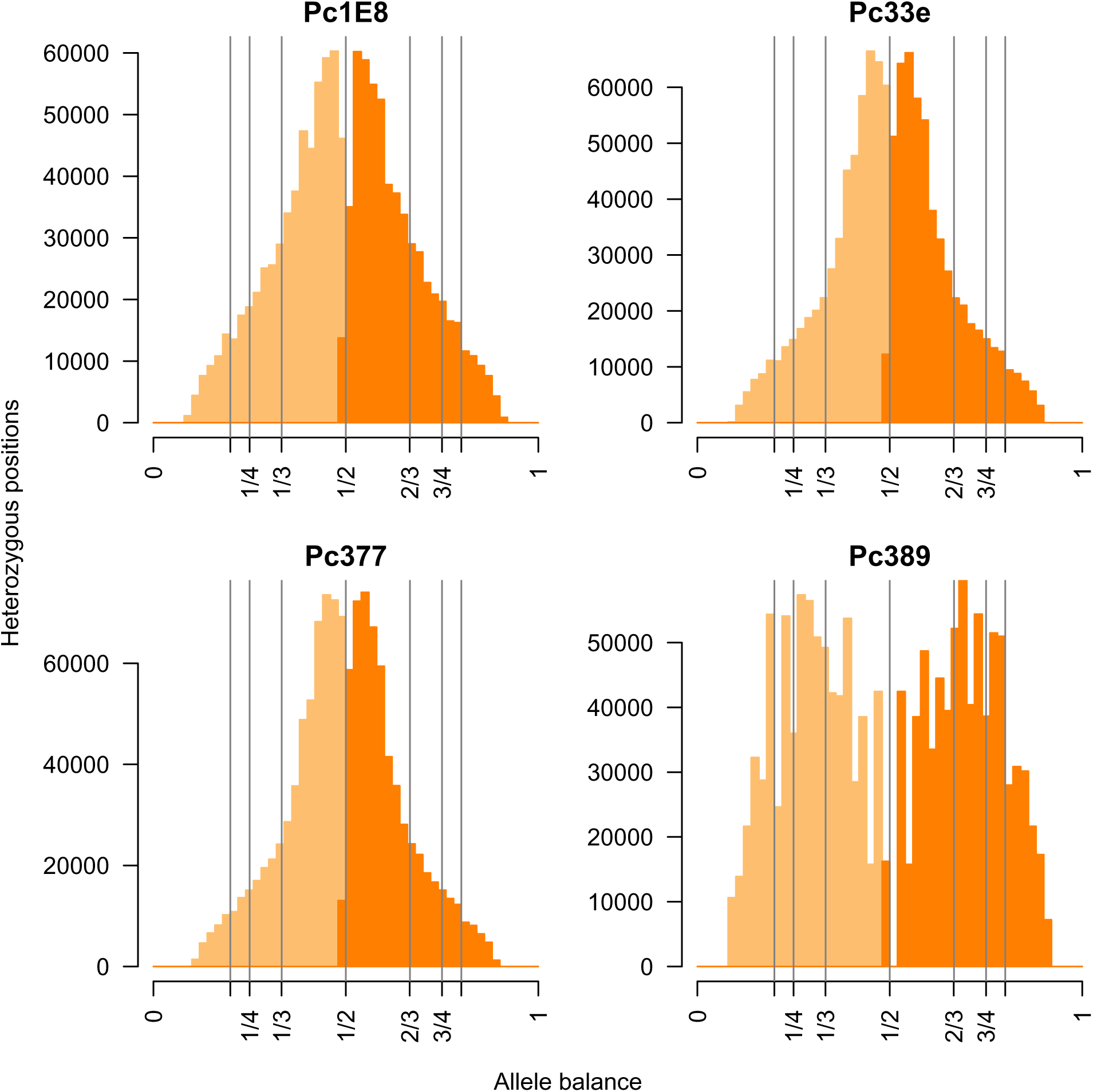
Histograms of allele balance for *P. capsici*. The expectations for pentaploid (1/5, 4/5), tetraploid (1/4, 3/4), triploid (1/3, 2/3), and diploid (1/2) heterozygote sequenced allele frequency is indicated with vertical lines.

**Figure Supplement 4B.**
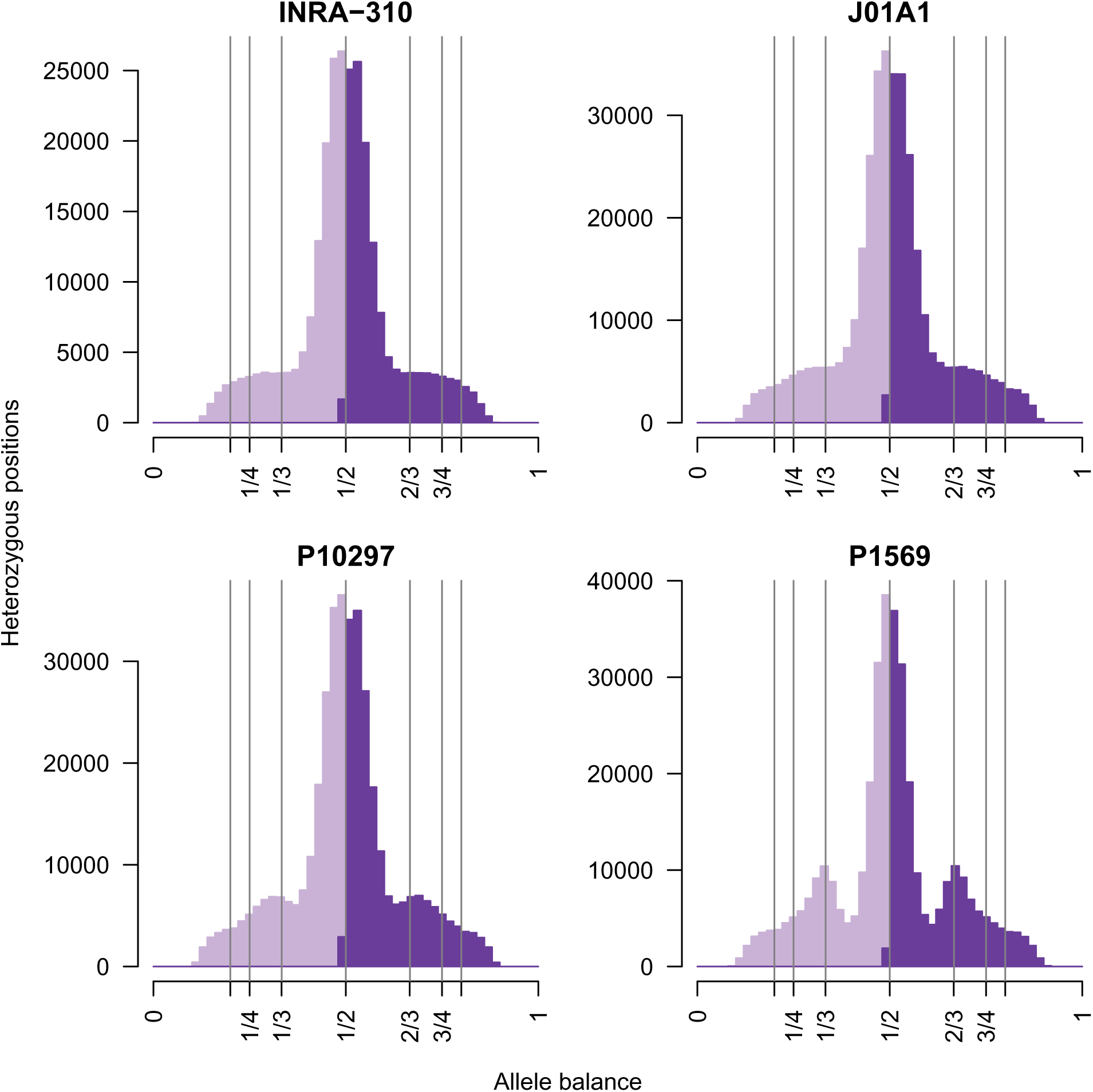
Histograms of allele balance for *P. parasitica*. The expectations for pentaploid (1/5, 4/5), tetraploid (1/4, 3/4), triploid (1/3, 2/3), and diploid (1/2) heterozygote sequenced allele frequency is indicated with vertical lines.

**Figure Supplement 4C.**
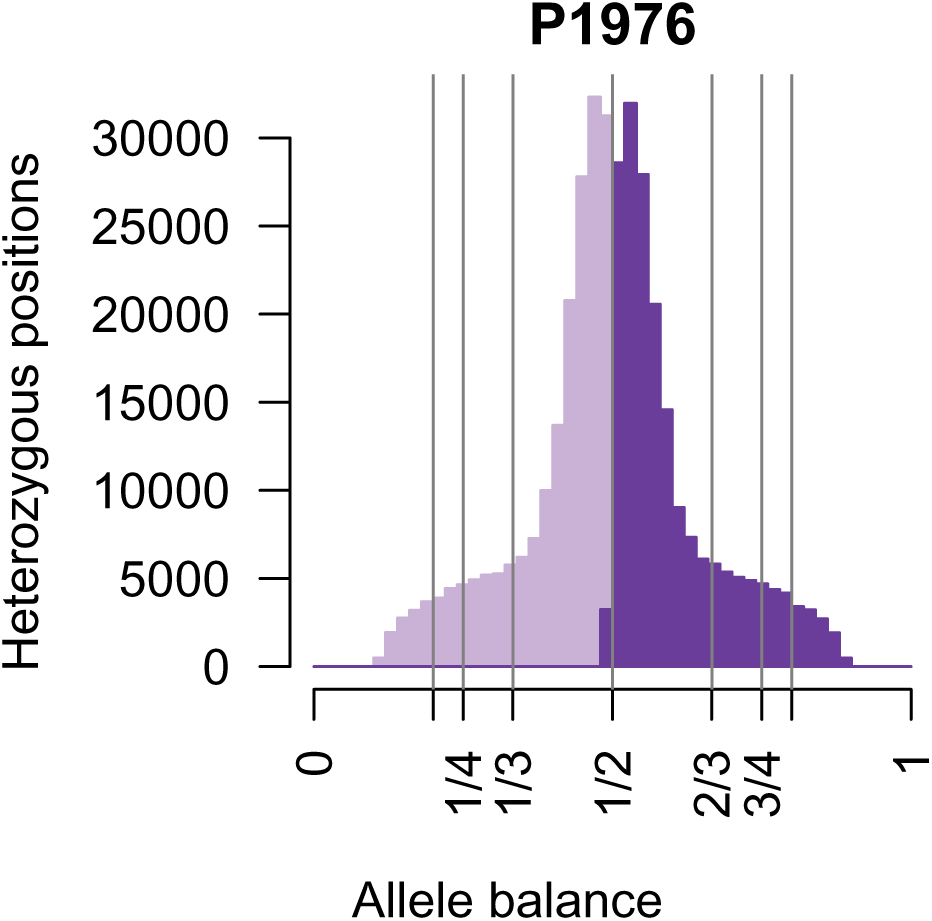
Histogram of allele balance for *P. parasitica*. The expectations for pentaploid (1/5, 4/5), tetraploid (1/4, 3/4), triploid (1/3, 2/3), and diploid (1/2) heterozygote sequenced allele frequency is indicated with vertical lines.

**Figure Supplement 4D.**
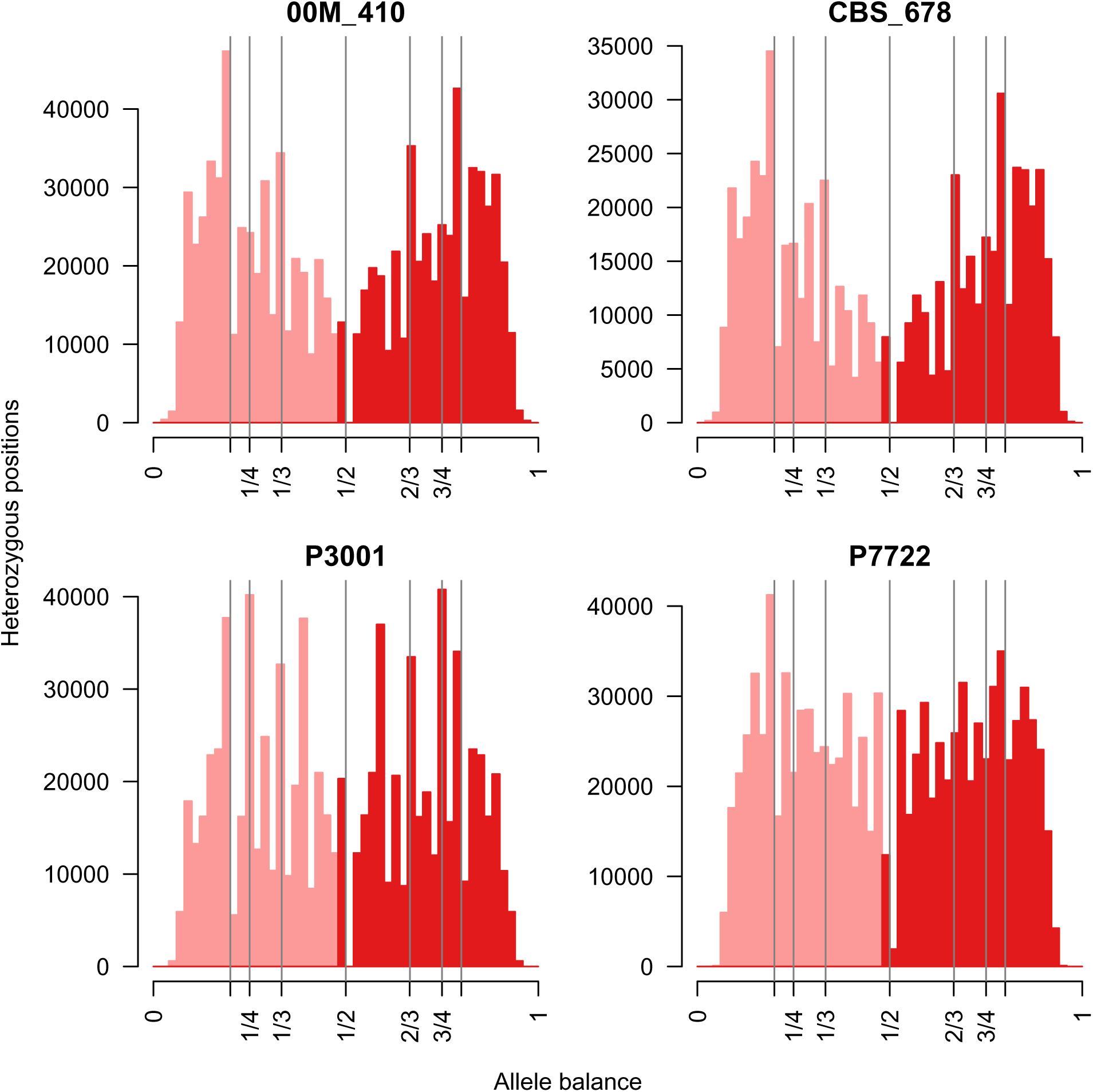
Histograms of allele balance for *P. mirabilis*. The expectations for pentaploid (1/5, 4/5), tetraploid (1/4, 3/4), triploid (1/3, 2/3), and diploid (1/2) heterozygote sequenced allele frequency is indicated with vertical lines.

**Figure Supplement 4E.**
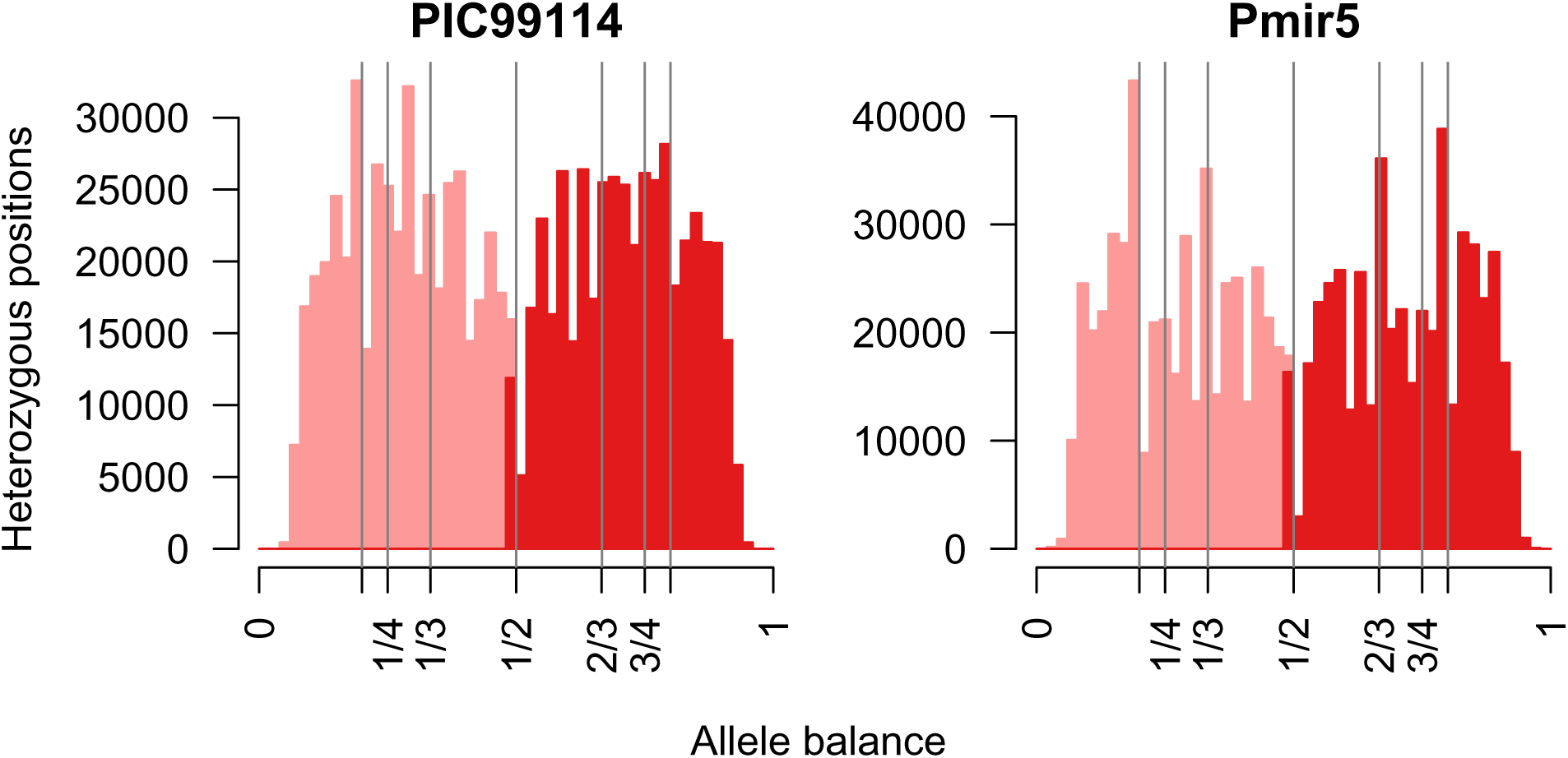
Histograms of allele balance for *P. mirabilis*. The expectations for pentaploid (1/5, 4/5), tetraploid (1/4, 3/4), triploid (1/3, 2/3), and diploid (1/2) heterozygote sequenced allele frequency is indicated with vertical lines.

**Figure Supplement 4F.**
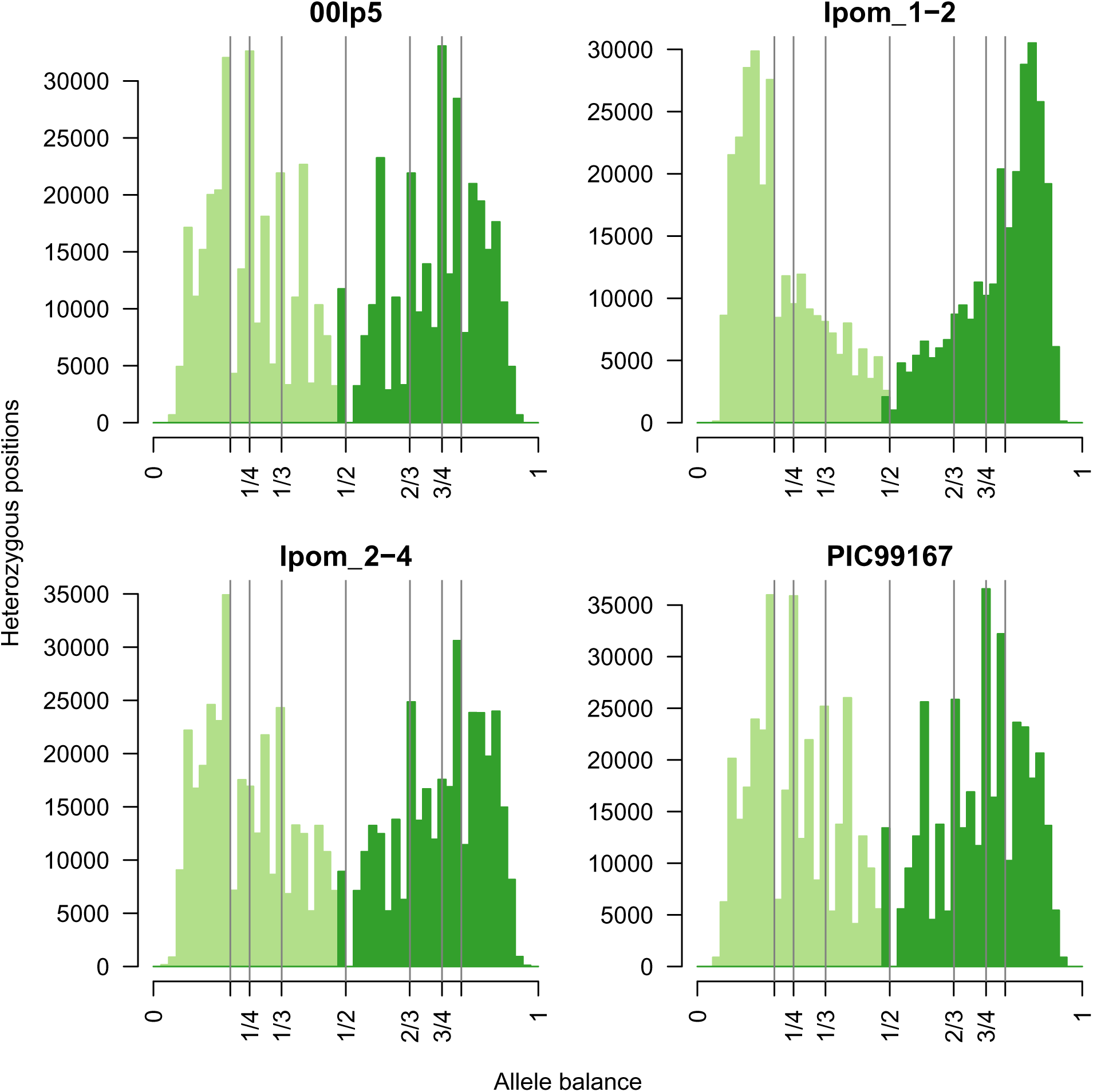
Histograms of allele balance for *P. ipomoeae*. The expectations for pentaploid (1/5, 4/5), tetraploid (1/4, 3/4), triploid (1/3, 2/3), and diploid (1/2) heterozygote sequenced allele frequency is indicated with vertical lines.

**Figure Supplement 4G.**
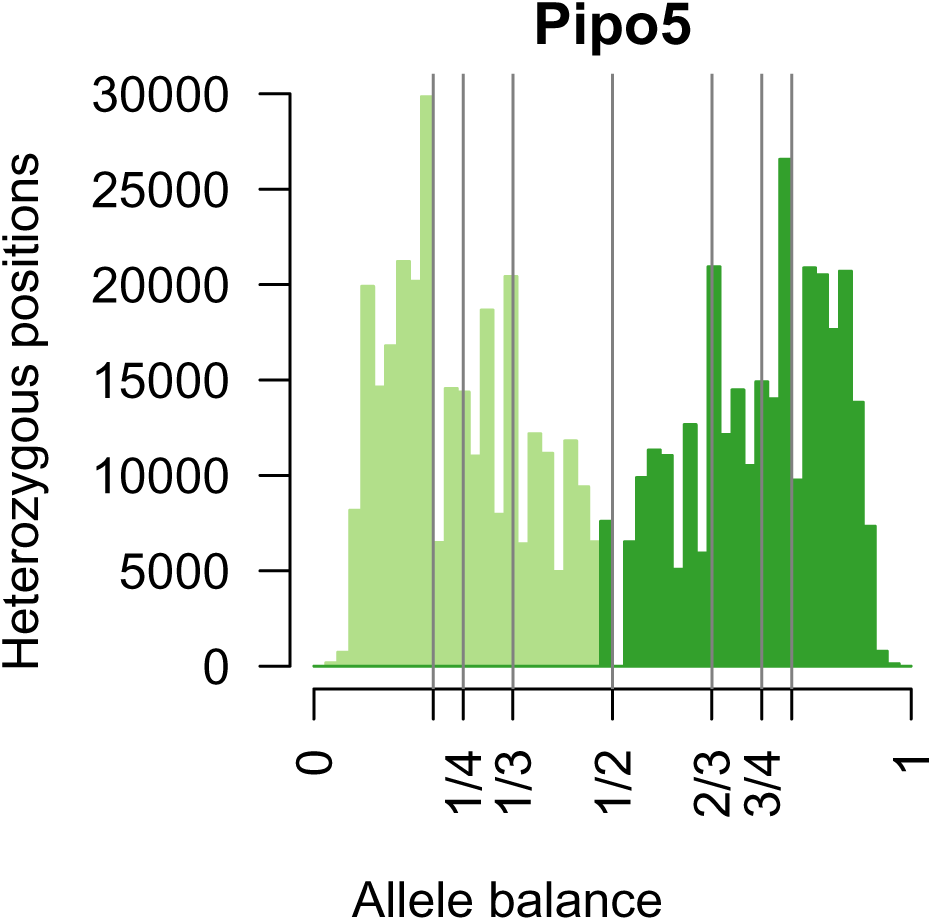
Histograms of allele balance for *P. ipomoeae*. The expectations for pentaploid (1/5, 4/5), tetraploid (1/4, 3/4), triploid (1/3, 2/3), and diploid (1/2) heterozygote sequenced allele frequency is indicated with vertical lines.

**Figure Supplement 4H.**
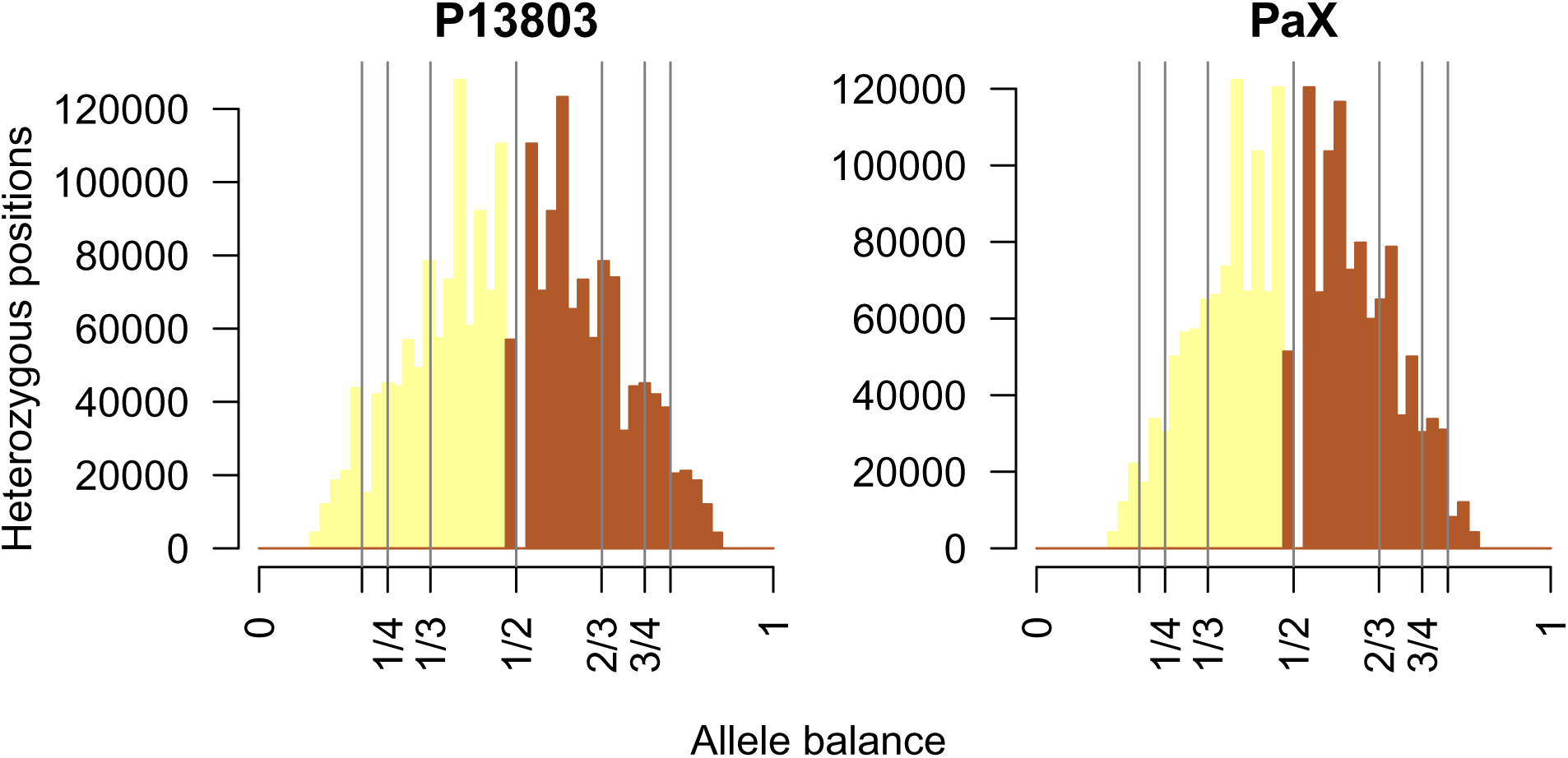
Histograms of allele balance for *P. andina*. The expectations for pentaploid (1/5, 4/5), tetraploid (1/4, 3/4), triploid (1/3, 2/3), and diploid (1/2) heterozygote sequenced allele frequency is indicated with vertical lines.

## 7 Gene loss categories

Gene loss, relative to T30-4, was more frequent in RxLR genes and was more frequent in Mexican isolates (sexually reproducing) than US-1 isolates (clonally reproducing). Core orthologous genes had among the lowest count of lost genes for Mexican, South American, and US-1 isolates. RxLR effector genes had the highest number of gene losses with Mexican isolates having a similar number of losses as South American isolates and US-1 isolates had a non-significantly lower number of gene losses.

**Figure Supplement 5.**
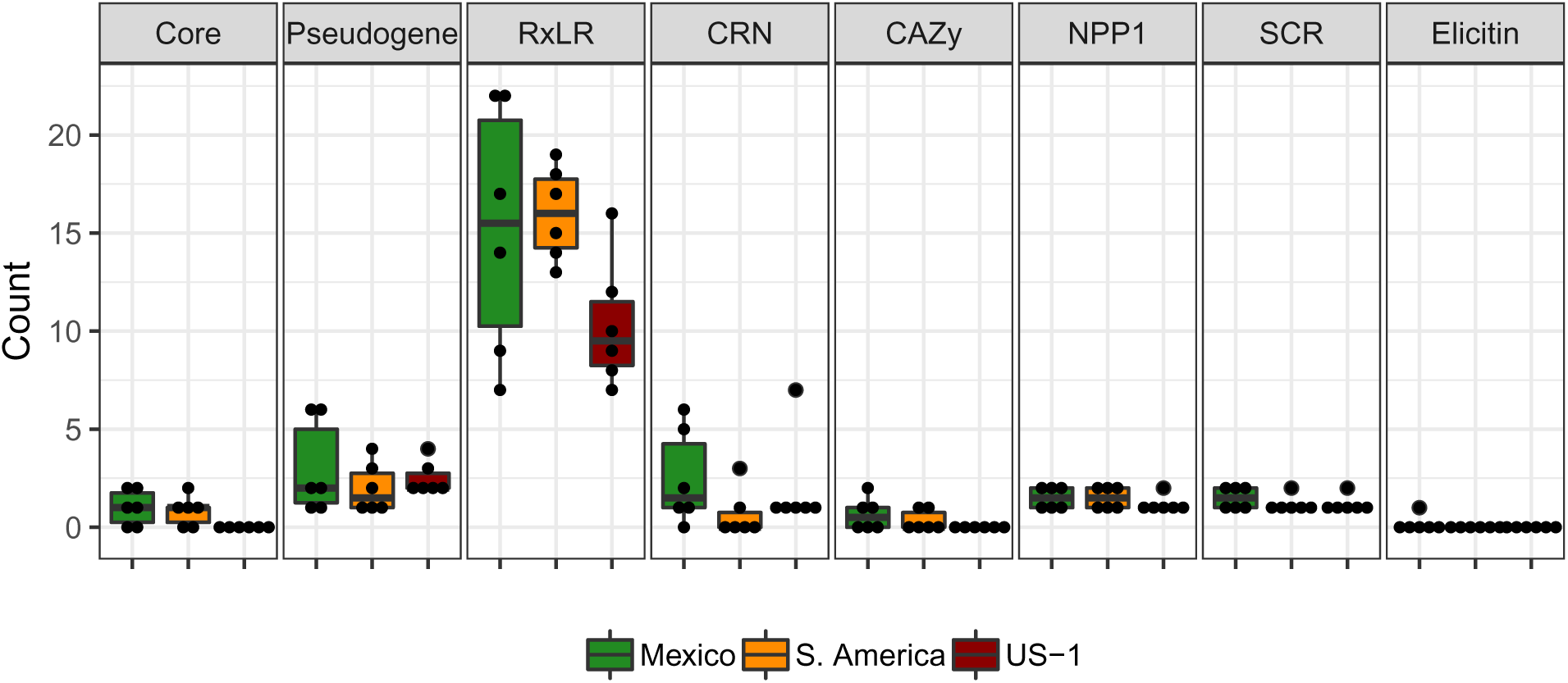
Number of gene losses observed in *P. infestans*, based on a breadth of coverage of zero, for annotated categories of genes.

